# Accuracy and data efficiency in deep learning models of protein expression

**DOI:** 10.1101/2021.11.18.468948

**Authors:** Evangelos-Marios Nikolados, Arin Wongprommoon, Oisin Mac Aodha, Guillaume Cambray, Diego A. Oyarzún

## Abstract

Many applications of synthetic biology involve engineering microbial strains to express high-value proteins. Thanks to advances in rapid DNA synthesis and sequencing, deep learning has emerged as a promising approach to build sequence-to-expression models for strain design and optimization. Such models, however, require large amounts of training data that are costly to acquire, which creates substantial entry barriers for many laboratories. Here, we study the relation between model accuracy and data efficiency in a large panel of machine learning models of varied complexity, from penalized linear regressors to deep neural networks. Our analysis is based on data from a large genotype-phenotype screen in *Escherichia coli*, which was generated with a design-of-experiments approach to balance coverage and depth of the genotypic space. We sampled these data to emulate scenarios with a limited number of DNA sequences for training, as commonly encountered in strain engineering applications. Our results suggest that classic, non-deep, models can achieve good prediction accuracy with much smaller datasets than previously thought, and provide robust evidence that convolutional neural networks further improve performance with the same amount of data. Using methods from Explainable AI and model benchmarking, we show that convolutional neural networks have an improved ability to discriminate between input sequences and extract sequence features that are highly predictive of protein expression. We moreover show that controlled sequence diversity leads to important gains in data efficiency, and validated this principle in a separate genotype-phenotype screen in *Saccharomyces cerevisiae.* These results provide practitioners with guidelines for designing experimental screens that strike a balance between cost and quality of training data, laying the groundwork for wider adoption of deep learning across the biotechnology sector.

## I. INTRODUCTION

Microbial production systems have found applications in many sectors of the economy^1^. In a typical microbial engineering pipeline, cellular hosts are transformed with heterologous genes that code for target protein products, and a key requirement is maximization of titers, productivity, and yield. Such optimization requires the design of genetic elements that ensure high transcriptional and translational efficiency^2^, such as promoter^3^ or ribosomal binding sequences^4^. However, prediction of protein expression is notoriously challenging and, as a result, strain development suffers from costly rounds of prototyping and characterization, typically relying on heuristic rules to navigate the sequence space towards increased production.

Progress in batch DNA synthesis and high-throughout sequencing has fueled the use of deep mutational scanning to study genotype-phenotype associations. Several works have combined high-throughput mutagenesis with a diverse range of measurable phenotypes, including protein expression^5–8^, ribosome loading^9^, and DNA methylation^10,11^. As a result, recent years have witnessed a substantial interest in machine learning methods that leverage such data for phenotypic prediction^9,12-15^. In synthetic biology, recent works have incorporated machine learning into the design-build-test cycle for predictive modelling of ribosomal binding sequences^16^, RNA constructs^17^, promoters^18^ and other regulatory elements^19^. Such sequence-to-expression models can be employed as *in silico* platforms for discovering variants with improved expression properties, paving the way toward a new level of computer-aided design of production strains^18^.

Deep learning algorithms, in particular, can uncover relations in the data on a scale that would be impossible by inspection alone, owing to their ability to capture complex dependencies with minimal prior assumptions^20^. Although deep learning models can produce highly accurate phenotypic predictions^12,21,22^, they come at the cost of enormous data requirements for training, typically ranging from tens to hundreds of thousands of sequences; see recent examples in Supplementary Table S1. Little attention has been paid to deep learning models in synthetic biology applications where data sizes are far below the requirements of state-of-the-art algorithms and, moreover, there is a poor grasp of what makes a good training dataset for model training. This is particularly relevant in applications where the cost of strain phenotyping is a limiting factor, as this places an upper ceiling on the number of variants that can be screened. The challenge is then to design a limited set of variants so that the resulting data can be employed to train useful predictors of protein expression. For example, if the sequence space has a broad and shallow coverage, i.e. composed of distant and isolated variants, the resulting data may be difficult to regress because each sample contains little information that correlates with expression. Conversely, if the coverage of the screen is narrow and deep, i.e. composed of closely related sequence variants, models may be accurate but generalize poorly to other regions of the sequence space.

Here, we trained a large number of sequence-to-expression models on datasets of variable size and sequence diversity. We employed a large screen of superfolder GFP-producing (sfGFP) strains in *Escherichia coli*^23^ that was designed to ensure a balanced coverage of the sequence space. We sampled these data so as to construct training datasets of varying size and controlled sequence diversity. We first establish the baseline performance of a range of classic, non-deep, machine learning models trained on small datasets with various phenotype distributions and using a range of strategies for encoding DNA sequences. This analysis revealed that for this particular dataset, accurate models can be trained on as few as a couple of thousand variants. We moreover show that convolutional neural networks (CNN), a common deep learning architecture, further improve predictions without the need to acquire further data. Using tools from Explainable AI^24^, we show that CNNs can better discriminate between input sequences than their non-deep counterparts and, moreover, the convolutional layers provide a mechanism to extract sequence features that are highly predictive of protein expression. We finally demonstrate that in limited data scenarios, controlled sequence diversity can improve data efficiency and improve predictive performance across larger regions of the sequence space. We validated this conclusion in a recent dataset of ~3,000 promoter sequences in *Saccharomyces cerevisiae*^25^. Our results provide a systematic characterization of sequence-to-expression machine learning models, with implications for the wider adoption of deep learning in strain design and optimization.

## II. RESULTS

### A. Size and diversity of training data

We sought to compare various machine learning models using datasets of different size and diversity. To this end, we employed the genotype-phenotype association data from Cambray *et al*^23^. This dataset comprises fluorescence measurements for an sfGFP-coding sequence in *Escherichia coli,* fused with more than 240,000 upstream 96nt regions that were designed to perturb translational efficiency and the resulting expression level. The library of upstream sequences was 1 randomized with a rigorous design-of-experiments approach so as to achieve a balanced coverage of the sequence space and a controlled diversity of variants. Specifically, 96nt sequences were designed from 56 seeds with maximal pairwise Hamming distances. Each seed was subject to controlled randomization using the D-Tailor framework^26^, so as to produce mutational series with controlled coverage of eight biophysical properties at various levels of granularity: nucleotide sequence, codon sequence, amino acid sequence, and secondary mRNA structure (Figure 1A).

**FIG. 1.**
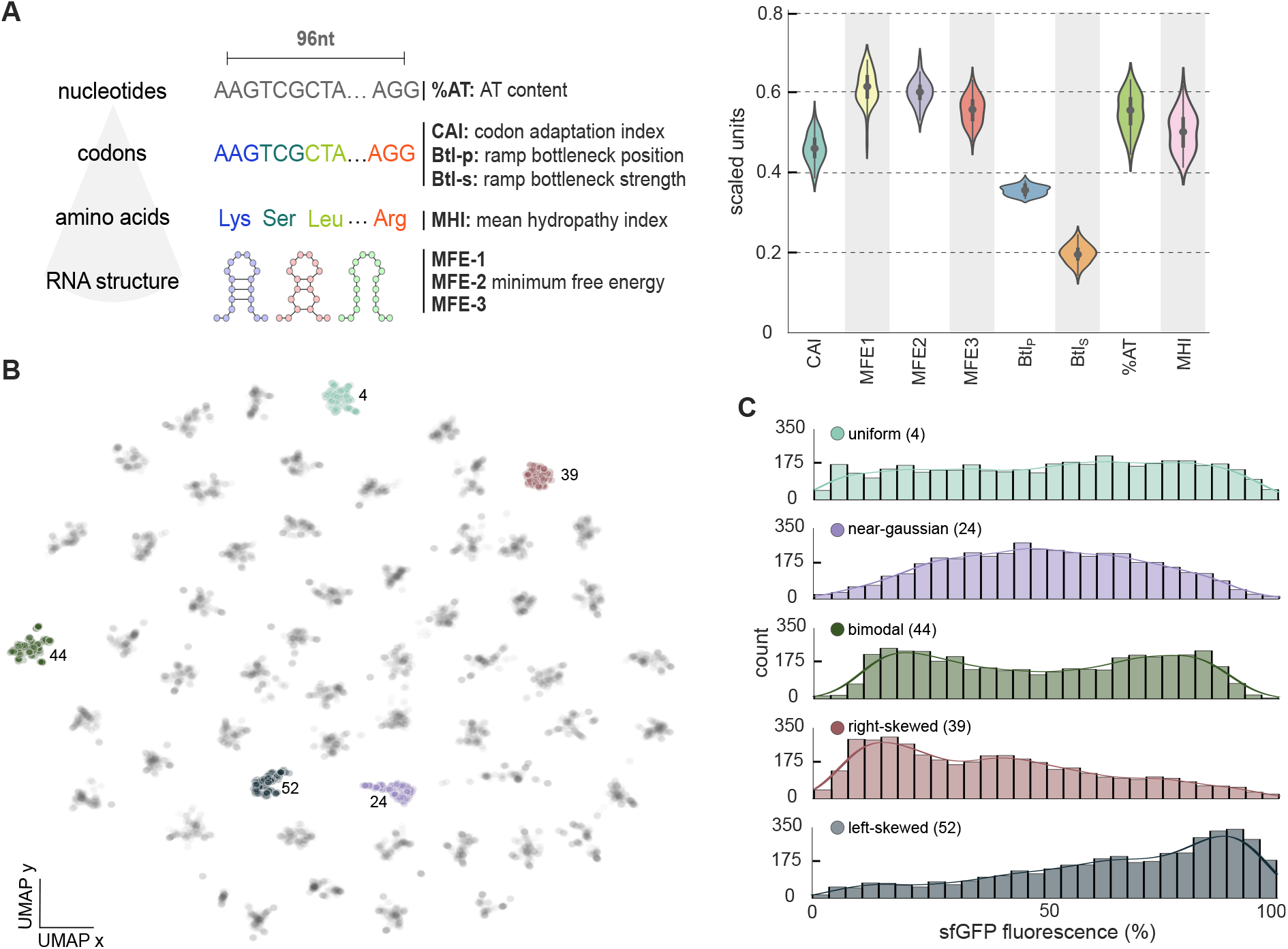
Characterization of the training data. **(A)** We employed a large phenotypic screen in *Escherichia coll^23^* of an sfGFP coding gene preceded by a variable 96nt sequence. The variable region was designed on the basis of eight sequence properties previously described as impacting translational efficiency: nucleotide content (%AT), patterns of codon usage (codon adaptation index, CAI, codon ramp bottleneck position, Btl_P_, and strength, Btl_S_), hydrophobicity of the polypeptide (mean hydrophobicity index, MHI) and stability of three secondary structures tiled along the transcript (MFE-1, MFE-2, and MFE-3). A total of 56 seed sequences were designed to provide a broad coverage of the sequence space, and then subjected to controlled randomization to create 56 mutational series of ~4,000 sequences each. After removal of variants with missing measurements, the dataset contains 228,000 sequences. Violin plots show the distribution of the average value of the eight properties across the 56 mutational series; the biophysical properties were normalized to the range [0,1] and then averaged across series. **(B)** Two dimensional UMAP^27^ visualization of overlapping 4-mers computed for all 228,000 sequences; this representation reveals 56 clusters, with each cluster corresponding to a mutational series that locally explores the sequence space around its seed; we have highlighted five series with markedly distinct phenotype distributions (labels denote the series number). Other UMAP projections for overlapping 3-mers and and 5-mers are shown in Supplementary Figure S1. **(C)** Mutational series with qualitatively distinct phenotypic distributions, as measured by FACS-sequencing of sfGFP fluorescence normalized to its maximal measured value; solid lines are Gaussian kernel density estimates of the fluorescence distribution. Measurements are normalized to the maximum sfGFP fluorescence across cells transformed with the same construct averaged over 4 experimental replicates of the whole library^23^. Fluorescence distributions for all mutational series are shown in Supplementary Figure S2.

The complete dataset contains 56 mutational series that provide wide coverage of the sequence space, while each series contains ~4,000 sequences for local exploration in the vicinity of the seed. The dataset is particularly well suited for our study because it provides access to controllable sequence diversity, as opposed to screens that consider either fully random sequences with limited coverage, or single mutational series that lack diversity.

To further characterize the sequence diversity across the library of 56 mutational series, we visualized the distribution of overlapping 4-mers using the Uniform Manifold Approximation and Projection (UMAP) algorithm for dimensionality reduction^27^. The resulting two-dimensional distribution of sequences (Figure 1B) shows a clear structure of 56 clusters, each corresponding to a mutational series. Moreover, the sfGFP fluorescence data (Figure 1C, Supplementary Figure S2) display marked qualitative differences across mutational series, including near-Gaussian distributions, left- and right-skewed distributions, as well as bimodal and uniform distributions. This indicates that the dataset is diverse in both genotype and phenotype space, and thus well suited for benchmarking machine learning models because it allows probing the impact of both genetic and phenotypic diversity on model accuracy.

### B. Impact of DNA encoding and sample size of training sequences

To understand the baseline performance of classic (non-deep) machine learning models, we trained various regressors on datasets of varying sizes and with different DNA encoding strategies I (Figure 2A). Sequence encoding is needed to featurize nucleotide strings into numerical vectors that can be processed by downstream machine learning models. We considered DNA encodings on three resolutions (Table I, Figure 2A): global biophysical properties (Figure 1A), DNA subsequences (overlapping *k*-mers), and single nucleotide resolution (one-hot encoding).

**FIG. 2.**
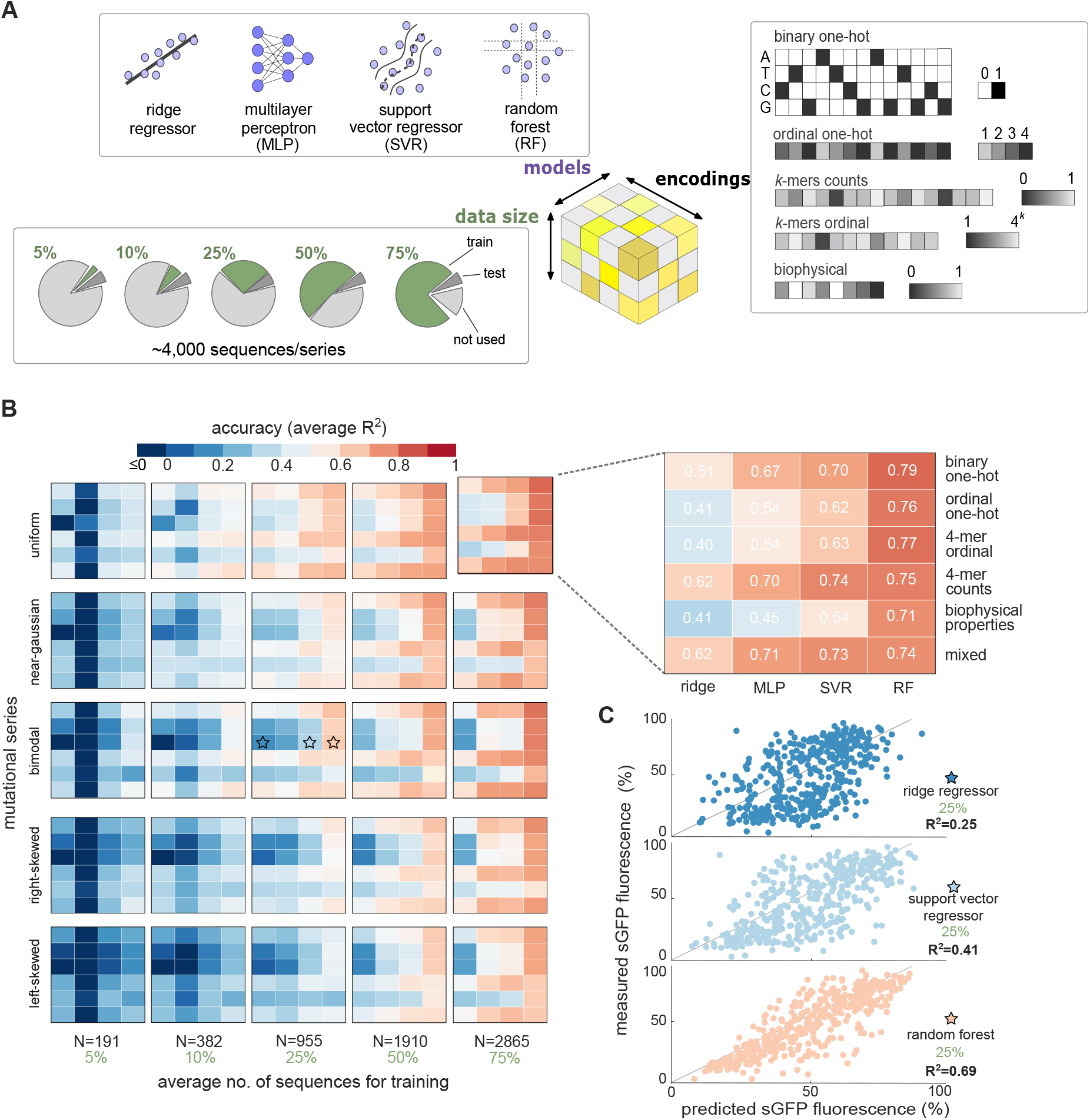
Accuracy of non-deep machine learning models. **(A)** We trained models using datasets of variable size and with different strategies for DNA encoding. Sequences were converted to numerical vectors with five DNA encoding strategies (Table I), plus an additional mixed encoding consisting of binary one-hot augmented with the biophysical properties of Figure 1A; in all cases, one-hot encoded matrices were flattened as vectors of dimension 384. We considered four non-deep models trained on an increasing number of sequences from five mutational series with different phenotype distributions (Figure 1B). **(B)** Impact of DNA encoding and data size on model accuracy. Overall we found that random forest regressors and binary one-hot encodings provide the best accuracy; we validated this optimal choice across the whole sequence space by training more than 5,000 models in all mutational series (Supplementary Figure S5). Phenotype distributions have a minor impact on model accuracy thanks to the use of stratified sampling for training. Model accuracy was quantified by the coefficient of determination (*R*^2^) between predicted and measured sfGFP fluorescence, computed on ~400 test sequences held-out from training and validation. The reported *R*^2^ values are averages across five training repeats with resampled training and test sets (Monte Carlo cross-validation). In each training repeat, we employed the same test set for all models and encodings. The full cross-validation results (Supplementary Figure S4) show robust performance and little overfitting, particularly for the best performing models. **(C)** Exemplar predictions on held-out sequences for three models from panel B (marked with stars); the shown models were trained on 25% of mutational series 44 (bimodal fluorescence distribution; Figure 1C) using 4-mer ordinal encoding. Details on model training and hyperparameter optimization can be found in the Methods, Supplementary Figure S3, and Supplementary Tables S2–S3.

**TABLE I.** DNA encodings for model training. We considered sequence encodings at three resolutions, which result in encoded vectors of different length. In the global encoding, sequences are described by the eight biophysical features employed in the original experimental design^23^(Figure 1A). At a subsequence resolution, we considered two versions of overlapping *k*-mers: an ordinal version where each *k*-mer is assigned a unique integer value between 1 and 4^*k*^, and *k*-mer counts containing the number of occurrences of each unique *k*-mer along the sequence; in our results we generally employed *k* = 4 for model training, but observed similar results for other choices of *k*. For base-resolution encodings, we employed two variants of one-hot encoding: binary one-hot where a sequence of length *L* is encoded as a binary matrix of size 4 ×L, with each column having a one at the position corresponding to the base in the sequence, and zeros elsewhere; ordinal one-hot encoding assigns a unique integer value to each of the four bases, resulting in encoded vectors of length L. Mixed encodings were constructed from flattened one-hot encoded matrices concatenated with the vector of biophysical properties, leading to feature vectors of dimension 4L + 8.

| DNA encoding | Resolution | Dimension | Positional |
| --- | --- | --- | --- |
| biophysical properties | global | 8 | × |
| $k$-mer counts | subsequence | $4^k$ | × |
| $k$-mer ordinal | subsequence | $L - k + 1$ | ✓ |
| binary one-hot | single base | $4L$ | ✓ |
| ordinal one-hot | single base | $L$ | ✓ |
| mixed | multiple | $4L + 8$ | ✓ |

We trained models on five mutational series chosen because of their markedly different expression distributions (Figure 1B), and with an increasing number of sequences for training (5%, 10%, 25%, 50% and 75% of sequences per series). Given the variation in phenotype distributions, we stratified training samples to ensure that their distribution is representative of the full series. We considered four non-deep models: ridge regressor^28^ (a type of penalized linear model), multilayer perceptrons^29^ (MLP, a shallow neural network with three hidden layers), support vector regressor^30^(SVR, based on linear separation of the feature space with a radial basis function kernel), and random forest regressor^31^ (RF, based on axis-aligned splits of the feature space). We chose this array of models because they markedly differ in their principle of operation and underlying assumptions on the shape of the feature space. We tuned model hyperparameters using grid search and 10-fold cross-validation on datasets assembled from aggregated fractions of all mutational series; this allowed us to determine a fixed set of hyperparameters for each of the four models with good performance across the whole dataset (see Methods and Supplementary Figure S3). In all cases, we assessed predictive accuracy using the coefficient of determination, *R*^2^ defined in Eq. (1), between measured and predicted sfGFP fluorescence computed on a set of ~400 test sequences (Supplementary Figure S3) that were held-out from model training and validation.

In line with expectation, the results in Figure 2B show that models trained on small datasets are generally poor irrespective of the encoding or regression method. Linear models (ridge) display exceptionally poor accuracy and are insensitive to the size of training set. In contrast, a shallow neural network (multilayer perceptron) achieved substantial gains in accuracy with larger training sets, possibly owing to its ability to capture nonlinear relationships. Our results show that mildly accurate models (*R*^2^ ≥ 50%) can be obtained from training sets with ~ 1,000 sequences using random forests and support vector regressors (Figure 2C). We found random forest regressors to be the most accurate among the considered models, consistently achieving *R*^2^ ≥ 50%for datasets with more than 1,000 samples and showing a stable performance when trained on other mutational series (Supplementary Figure S5). To produce robust performance metrics, the *R*^2^ scores in Figure 2B are averages across five training repeats with resampled training and test sets (Monte Carlo cross-validation).

We also observed a sizeable impact of DNA encodings on prediction accuracy. Subsequence-resolution encodings achieve varying accuracy that is highly dependent on the specific mutational series and chosen model (Figure 2B, Supplementary Figure S5). Overall we found a strong preference for base-resolution encodings, with binary one-hot representations achieving the best accuracy. A salient result is that the sequence biophysical properties led to poorer accuracy than most other encodings, possibly due to their inability to describe a high-dimensional sequence space with a relatively small number of features (8). Their poor performance is particularly surprising because the biophysical properties were used to design the sequences based on their presumed phenotypic impact^23^; moreover, some of them (codon adaptation index, mRNA secondary structures) represent the state-of-the-art understanding of a sequence impact on translation efficiency^23,32,33^, while the best performing one-hot encodings lack such mechanistic information. In an attempt to combine the best of both approaches, we trained models on binary one-hot sequences augmented with the biophysical properties (“mixed” encoding in Table I, Figure 2B and Supplementary Figure S5). This strategy led to slight gains in accuracy for small training sets; e.g. for ~200 training 1 sequences, the median *R*^2^ with mixed encoding is 0.30 vs a median of 0.26 for binary one-hot (Supplementary Figure S5). For larger training sets, however, binary one-hot encodings gave the best and most robust accuracy across models.

### C. Deep learning improves accuracy without more data

Prior work has shown that deep learning can produce much more accurate predictions than non-deep models^16,19^. Deep learning models, however, typically require extremely large datasets for training; some of the most powerful deep learning phenotypic predictors, such as DeepBIND^12^, Optimus 5-prime^9^, ExpressionGAN^34^, and Enformer^14^ were trained with tens to hundreds of thousands of variants. In the case of sequence-to-expression models, recent literature shows a trend towards more complex and data-intensive models (see Supplementary Table S1); the most recent sequence-to-expression model employed ~20,000,000 promoter sequences to predict protein expression in *Saccharomyces cerevisiae*^25^. It is often unclear if the accuracy of such deep learning predictors results from the chosen model architecture or simply from the sheer size of the training data. To test this idea with our data, we designed a convolutional neural network (CNN, a common type of deep learning model) with an off-the-shelf architecture of similar complexity to those employed in recent literature^25^.

Our CNN architecture (Figure 3A) processes a binary one-hot encoded sequence through three convolutional layers, followed by a four dense layers which are equivalent to a four-layer multilayer perceptron. The convolutional layers can be regarded as weight matrices acting on an input sequence. By stacking several convolutional layers, the network can capture interactions between different components of the input. We designed the CNN architecture with a Bayesian optimization algorithm^35^ to determine the optimal number of network layers, as well as the optimal settings for the filters in each layer (see Supplementary Tables S4–S5 for details). In addition to the components shown in Figure 3A, we also included a dropout layer to prevent overfitting and max pooling to reduce the number of trainable parameters. Similar as with the non-deep models in Figure 2B, hyperparameter optimization was performed by splitting the data into separate sets for training and cross-validation (details in Methods and Supplementary Figure S3). This allowed us to find a single CNN architecture with good performance across the individual 56 mutational series and the whole dataset.

**FIG. 3.**
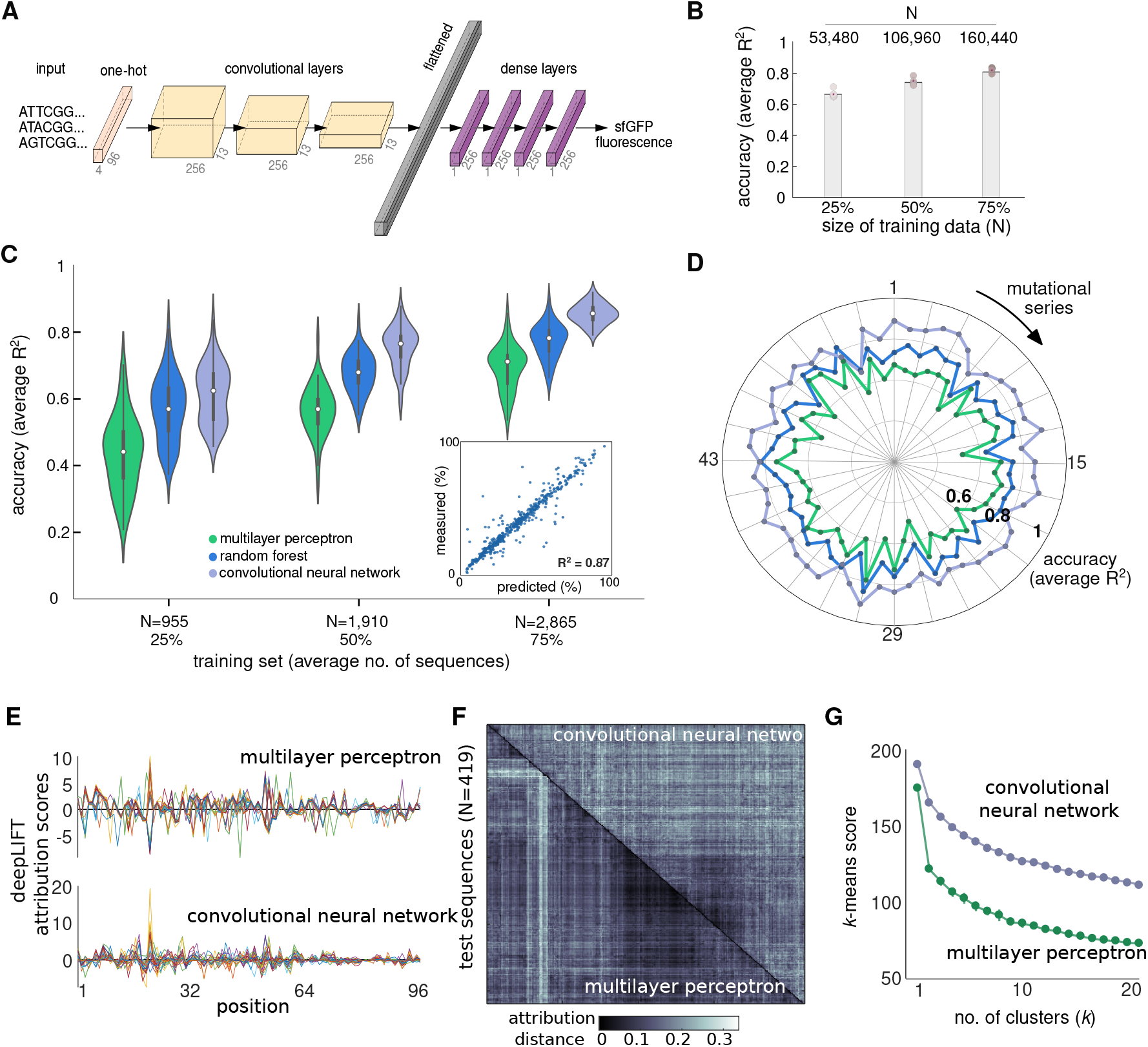
Prediction accuracy of deep neural networks. **(A)** Architecture of the convolutional neural network (CNN) employed in this paper; the output is the predicted sfGFP fluorescence in relative units. The CNN architecture was designed with Bayesian optimization^35^ to find a single architecture for all mutational series; our strategy for hyperparameter optimization can be found in the Methods, Supplementary Figure S3, and Supplementary Tables S4–S5. **(B)** Accuracy of the CNN in panel A trained on all mutational series. *R*^2^ values were computed on held-out sequences (10% of total) and averaged across 5 training repeats; bars denote the mean **R*^2^*. (**C**) Prediction accuracy of CNNs against random forest (RF) and multilayer perceptrons (MLPs) on all 56 mutational series using binary one-hot encoding. The CNNs yield more accurate predictions with the same training data. Violin plots show the distribution of 56 *R*^2^ values for each model averaged across 5 training repeats; *R*^2^ values were computed on held-out sequences (10% of sequences per series). Inset shows predictions of a CNN trained on 75%of the mutational series with a right-skewed phenotypic distribution (Figure 1B) computed on held-out test sequences. The CNNs are more complex than the shallow MLPs (2,702,337 *vs* 58,801 trainable parameters, respectively), but we also found that the CNNs outperform MLPs of comparable complexity (Supplementary Figure S8); this suggests that improved performance is a result of the convolutional layers acting as a feature extraction mechanism. Details on CNN training can be found in the Methods and Supplementary Figure S7. **(D)** Average *R*^2^ scores for each model across all 56 mutational series using 75% of sequences for training. (**E**) DeepLIFT^37^ attribution scores per nucleotide position for a given test sequence and trained model. Panels show scores of 30 sequences chosen at random from the same test set employed in C for models trained on 75% of mutational series 21. (**F**) Attribution distances for models trained on series 21. We computed the cosine distance between DeepLIFT scores for each sequence in the test set. Distance heatmaps were hierarchically clustered to highlight the cluster structure that both models assign to the input sequences. (**G**) *K*-means clustering of the distance matrices in panel E. Line plots show the optimal *k*-means score averaged across 20 runs with random initial cluster assignments. Lower scores for all values of *k* suggests that the MLP clusters sequences more heavily than the CNN; we found this pattern in all but four mutational series (Figure S10).

When trained on up to 75% of the full dataset (~160,000 sequences), our CNN model produced excellent predictions in test sets covering broad regions of the sequence space (average *R*^2^= 0.82 across five cross-validation runs, Figure 3B and Supplementary Figure S6). This suggests data size alone is sufficient for training accurate regressors, but our concern is that data of such scale are rarely available in synthetic biology applications. We thus sought to determine the capacity of CNNs to produce accurate predictions from much smaller datasets than previously considered in the literature. To this end, we trained CNNs with the same architecture in Figure 3A on each mutational series, using ~1,000–3,000 sequences in each case; details on CNN training can be found in the Methods, Supplementary Figure S7, and Supplementary Tables S4–S5. We benchmarked the accuracy of the CNNs against non-deep models trained on the same 56 mutational series. As benchmarks we chose two non-deep models: a shallow perceptron because it is also 1 a type of neural network, and a random forest regressor because it showed the best performance so far (Figure 2B). We found that CNNs are consistently more accurate than non-deep models, regardless of the size of the training data (Figure 3C-D) and across most of the 56 mutational series. In fact, in more than half of mutational series, the CNNs achieve accuracy over 60% with I ~1000 training sequences, and in some cases they reach near state-of-the-art accuracy (*R*^2^= 0.87 averaged across five cross-validation runs, Figure 3C inset). When trained on ~3,000 sequences, the CNNs outperformed the MLP in all mutational series, and the random forest regressor in all but four series (Figure 3D).

To understand how CNNs can provide such improved accuracy without larger training data, we compared them against multilayer perceptrons (MLPs) of increasing depth. We note that the CNN 1 in Figure 3C has ~45-fold more trainable parameters than the MLPs, which suggests that such additional complexity may be responsible for the improved predictive accuracy. We thus sought to determine if increasing MLP complexity could bring their performance to a level comparable to the CNNs. We trained deep MLPs with an increasing number of hidden layers on ~3,000 sequences from each mutational series. We found that the additional layers provide marginal improvements in accuracy, and that the performance gap between CNNs and MLPs exists even when both have a comparable number of trainable parameters (Supplementary Figure S8). This suggests that the higher accuracy of the convolutional network stems from its inbuilt inductive bias that enables it to capture local structure via the learned filters and more global structure through successive convolutional layers^36^. As a result, it can capture interactions between different components of the 1 input and produce sequence embeddings that are highly predictive of protein expression.

To further determine how the models process the input sequences, we employed methods from Explainable AI to quantify the sensitivity of both neural network models (shallow MLPs and CNNs) to changes in the input sequence. We utilized DeepLIFT^37^, a computationally efficient method that produces importance scores for each feature of the input; such scores are known as “attribution scores” in the Explanable AI literature^24^. When applied to one-hot encoded sequences, DeepLIFT produces scores at the resolution of single nucleotides (Figure 3E). We employed these scores to compute pairwise distances between sequences processed by the same model. The shorter that distance, the more the two sequences are detected as similar by the model. We computed such distances for all pairs of sequences in each test set processed by the MLP or CNN. The matrices of pairwise distances (Figure 3F) were then subjected to hierarchical clustering as a means to contrast the diversity of responses elicited by test sequences on the two models. Using *k*-means clustering, we showed that the CNN produces less clustered attribution distances than the MLP (Figure 3G), thus highlighting the ability of the CNN to discriminate input sequences with finer granularity than the MLP. This trend was found in all but four of the CNNs (Supplementary Figure S10).

### D. Impact of sequence diversity on model coverage

A well recognized caveat of sequence-to-expression models is their limited ability to produce accurate predictions in regions of the sequence space not covered by the training data^25,38^; this is commonly referred to as *generalization performance* in the machine learning jargon. In line with expectation, we found that the CNNs from Figure 3C, which were trained on a single mutational series each, performed poorly when tested on other mutational series (*R*^2^ ≤ 0 for most models, Supplementary Figure S11A); we observed similarly poor results for the non-deep models in Figure 2 (Supplementary Figure S11B). Negative *R*^2^ scores indicate an inadequate model structure with a poorer fit than a baseline model that simply predicts the average observed fluorescence. This means that models trained on a particular region of the sequence space are too specialized, and their phenotypic predictions do not generalize to distant sequences. Although poor generalization can be caused by model overfitting, our cross-validation results (see Supplementary Figure S6A and Supplementary Figure S7) rule out this option and suggest that it is rather a consequence of the large genotypic differences between mutational series, compounded with the high-dimensionality of the sequence space.

Recent work by Vaishnav and colleagues demonstrated that model generalization can be improved with CNNs of similar complexity to ours^25^ trained on extremely large data (~20,000,00 variants). Since the cost of such large screens is prohibitive in most synthetic biology applications, we sought to understand how model coverage could be improved in scenarios where data size is strongly limited. The idea is to design a sequence space for training that can enlarge the high-confidence regions of the predictors with a modest number of variants; this is somewhat akin to the concept of “informed training sets” recently introduced in the context of protein design^39^. To this end, we performed a computational experiment designed to test the impact of sequence diversity on the ability of CNNs to produce accurate predictions across different mutational series.

We trained CNNs on datasets of constant size but increasing sequence diversity (Figure 4A, Supplementary Figure S12). We considered an initial model trained on 5,800 sequences sampled from the aggregate of two series chosen at random, e.g. 2,900 sequences from series 13 and 23, respectively (Figure 4A top row). We successively added two series to the aggregate and retrained a CNN while keeping a constant number of total sequences. This results in sparser sampling from each mutational series and an increasingly diverse training set. For example, the second model (Figure 4A) was trained on 1,450 sequences from series 13, 23, 48 and 55, respectively. Overall, we trained a total of 27 models, the last of which comprises as few as 107 sequences per mutational series. The resulting models display substantial variations in their predictive power (Figure 4A). Most models displayed variable *R*^2^ scores across different series, and we identified two salient patterns: some series that are consistently well predicted even in small data scenarios (e.g. series 31 and 51), and some series are particularly hard to regress (e.g. series 28 and 54), which possible require a bespoke CNN architecture different from the one in Figure 3A. The results also show that increased diversity has a minor impact on model generalization; although some series not included in training do have improved prediction scores (e.g. series 53 in Figure 4A), we suspect this is likely a result of series being particularly easy to regress. In general, we observed patterns of low or negative *R*^2^ scores for series not included in the aggregate. Similar results were observed for other random choices of mutational series employed for training (Supplementary Figure S12).

**FIG. 4.**
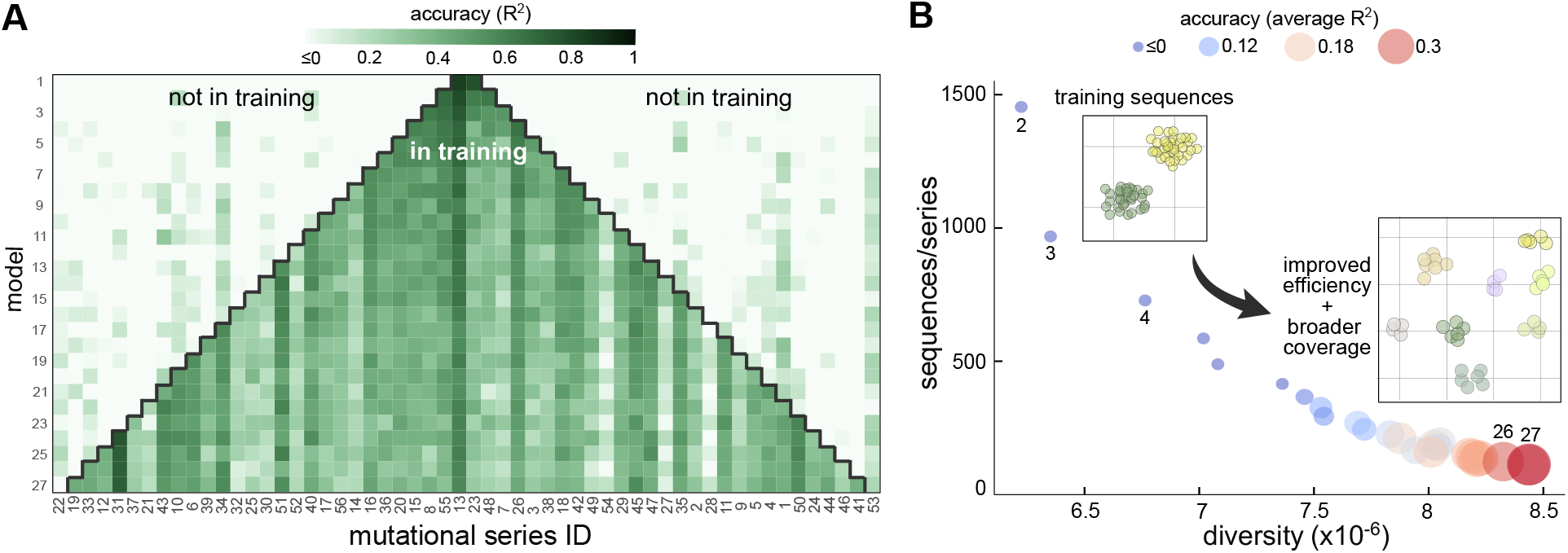
Impact of sequence diversity on model coverage. (**A**) We trained CNNs on datasets of constant size and increasing sequence diversity. We trained a total of 27 models by successively aggregating fractions of randomly chosen mutational series into a new dataset for training; the total size of the training was kept constant at 5,800 sequences. Training on aggregated sequences achieves good accuracy for mutational series in the training set, but poor predictions for series not included in the training data. This suggests that CNNs generalize poorly across unseen regions of the sequence space. Accuracy is reported as the *R*^2^ computed on 10% held-out sequences from each mutational series. We excluded two series from training to test the generalization performance of the last model. (**B**) Bubble plot shows the *R*^2^ values averaged across all mutational series for each model. Labels indicate the model number from panel A, and insets show schematics of the sequence space employed for training; for clarity, we have omitted model 1 from the plot. Improved sequence diversity leads to gains in predictive accuracy across larger regions of the sequence space; we observed similar trends for other random choices of series included in the training set (Supplementary Figure S12). The decreasing number of training sequences per series reflects better data efficiency, thanks to an increasingly diverse set of training sequences. To quantify sequence diversity, we counted the occurrence of unique overlapping 5-mers across all sequences of each training set, and defined diversity as 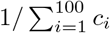 where *c_i_* is the count of the *i*-th most frequent 5-mers.

Crucially, the results in Figure 4B suggest that increased sequence diversity enlarges the region where the CNN can produce accurate predictions without increasing the size of the training data. We found that *R*^2^ > 30%in many regions of the sequence space can be achieved by models trained on just over a hundred sequences from those regions (e.g. model 27 in Figure 4A). For comparison, the CNN trained on all series without controlled diversity can double that accuracy, but with a 9fold increase in the size of the training data (*R*^2^= 0.65 for *N*=53,480 in Figure 3B). This means that model coverage can be enlarged with shallow sampling of previously unseen regions of the sequence space, which provides a useful guideline for experimental design of screens aimed at training sequence-to-expression models on a limited number of variants.

To test the validity of this principle in a different expression chassis and construct library, we repeated the computational experiment in Figure 4 using a recent genotype-phenotype screen of promoter sequences in *Saccharomyces cerevisiae*^25^. These data are comparable to the screen in Cambray et al^23^ in the sequence length (80nt) and its highly clustered coverage of genotypic space (Figure 5A). This clustered structure results from the design of the library itself, which is composed of 3,929 variants of 199 natural promoters. A key difference between this new dataset and Cambray et al^23^ is the construct architecture; unlike the UTR sequences in Figure 1B, promoter sequences account for regulatory effects but do not undergo transcription. Akin to our results in Figure 4, we aimed at testing the accuracy of machine learning regressors trained on datasets of constant size but increasing sequence diversity. Since this dataset contains a small number of variants for each gene (on average 20 variants/gene, see inset of Figure 5A), we first randomly aggregated the variant clusters into twelve groups containing an average of 327 sequences/group. We subsequently trained five Random Forest models on *N =* 400 binary one-hot encoded sequences drawn from different groups. For example, as shown in the Figure 5B, model 1 was trained on 200 sequences from two groups, whereas model 2 was trained on 100 variants from four groups. The training results (Figure 5B) show a strikingly similar pattern to those observed in our original dataset in Figure 4, thus strongly suggesting that sequence diversity can be exploited to train models with broader coverage and improved data efficiency.

**FIG. 5.**
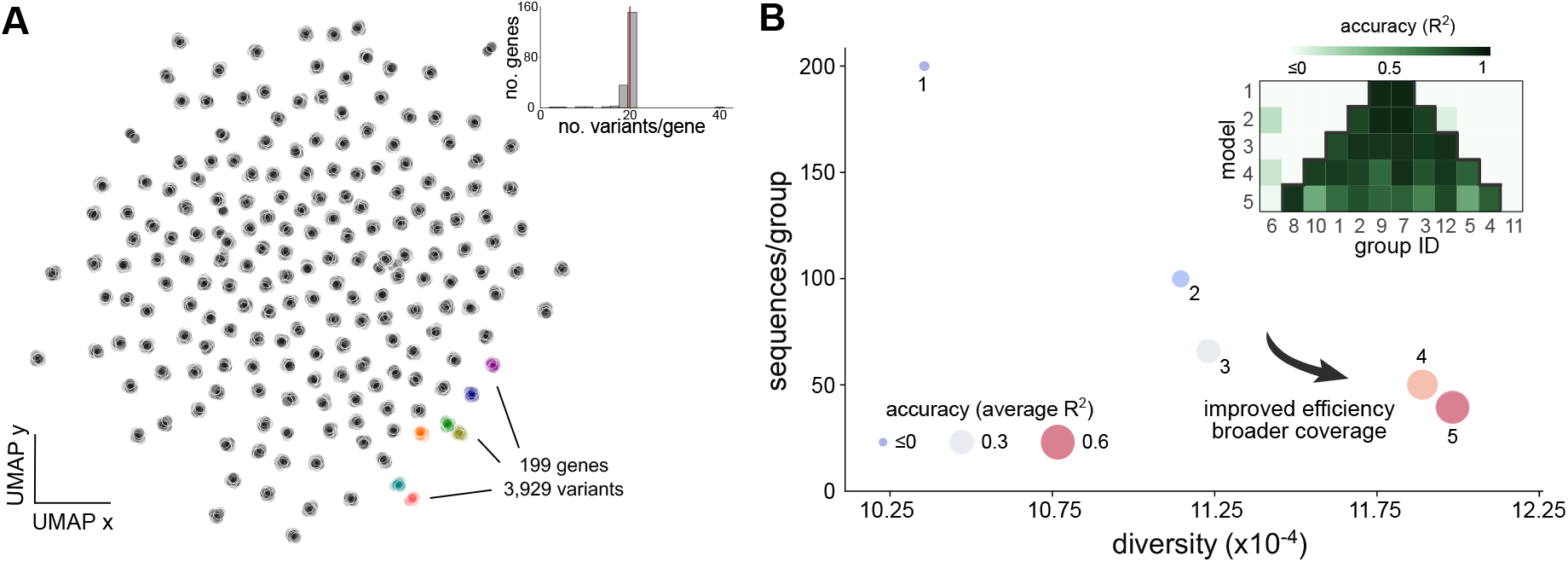
Sequence-to-expression models using promoter data from *Saccharomyces cerevisiae*. (**A**) Genotypic space of yeast promoter data from Vaishnav et al^25^ visualized with the UMAP^27^ algorithm for dimensionality reduction; sequences were featurized using counts of overlapping 4-mers, as in Figure 1B. The dataset contains 3,929 promoter variants (80nt long) of 199 native genes, as well as fluorescence measurements of a yellow fluorescent protein (YFP) reporter; inset shows the distribution of variants per gene across the whole dataset. **(B)** Bubble plots show the accuracy of five random forest (RF) models trained on datasets of constant size and increasing sequence diversity, following a similar strategy as in Figure 4A. We first aggregated variant clusters into twelve groups, and then trained RF models by aggregating fractions of randomly chosen groups into a new dataset for training; the total size of the training set was kept constant at 400 sequences. Accuracy was quantified with the *R*^2^ score averaged across test sets from each group (~30 sequences/group) that were held out from training. Inset shows model accuracy in each test set. In line with the results in Figure 4A, we observe that model coverage can be improved by adding small fractions of each group into the training set; we observed similar trends for other random choices of groups included in the training set (Supplementary Figure S13). Details on data processing and model training can be found in the Methods and Supplementary Text. Sequence diversity was quantified as in Figure 4B.

## III. DISCUSSION

Progress in high-throughput methods has led to large improvements in the size and coverage of genotype-phenotype screens, fuelling an increased interest in deep learning algorithms for phenotypic prediction^9,12-14,16,17,19,34^. Synthetic biology offers a host of applications that would benefit from such predictors, e.g. for optimization of protein-producing strains^40^, selection of enzymatic genes in metabolic engineering^41^, or the design of biosensors^42^. An often-overlooked limitation is that deep learning models require huge amounts of data for training, and the sheer cost of the associated experimental work is a significant barrier for most laboratories. Recent sequence-to-expression models have focused primarily on datasets with tens to hundreds of thousands of training sequences (Supplementary Table S1). While large data requirements are to be expected for prediction from long sequences such as entire protein coding regions, synthetic biologists often work with much shorter sequences to control protein expression levels (e.g. promoters^3^, riboso-mal binding sequences^4^, terminators^43^ and others). From a machine learning standpoint, shorter sequences offer potential for training models with smaller datasets, which can lower the entry barriers for practitioners to adopt deep learning for strain optimization.

Here, we examined a large panel of machine learning models, with particular emphasis on the relation between prediction accuracy and data efficiency. We used data from an experimental screen in which sequence features were manipulated using a Design of Experiments approach to perturb the translation efficiency of an sfGFP reporter in *E. coli*^23^. Thousands of local mutations were derived from more than fifty sequence seeds, yielding mutational series that enable deep focal coverage in distinct areas of the sequence space (Figure 1B). By suitable sampling of these data, we studied the impact of the size and diversity of training sequences on the quality of the resulting machine learning models.

Our analysis revealed two key results that can help incentivize the adoption of machine and deep learning in strain engineering. First, in our dataset we found that the number of training sequences required for accurate prediction is much smaller than what has been shown in the literature so far^8^·^12^·^16^·^17^·^25^. Traditional non-deep models can achieve good accuracy with as few as 1,000–2,000 sequences for training (Figure 2B). We moreover showed that deep learning models can further improve accuracy with the same amount of data. For example, our convolutional neural networks achieved gains of up to 10% in median prediction scores across all mutational series when trained on the same 2,000 sequences as the non-deep models (Figure 3C). Such performance improvement is a conservative lower bound, because we employed a fixed network architecture for all mutational series; further gains in accuracy can be obtained with custom architectures for different mutational series.

Second, we found that sequence diversity can be exploited to increase data efficiency and enlarge the sequence space where models produce reliable predictions. Using two different datasets with a similar structure of their sequence coverage, the *E. coli* library from Cambray et al^23^ as well as a recently published library of *S. cerevisiae* promoters^25^, we showed that machine learning models can expand their predictions to entirely new regions of the sequence space by training on a few additional samples from that region (Figures 5). This means that controlled sequence diversity can improve the coverage of sequence-to-expression models without the need for more training data. In other words, instead of utilizing fully randomized libraries for training^8,16,18^, it may be beneficial to first design few isolated variants for coverage, and then increase the depth with many local variants in the vicinity of each seed. Our work strongly suggests that such balance between coverage and depth can be advantageous in small data scenarios, where fully randomized libraries would lead to datasets with faraway and isolated sequences that inherently require large datasets to achieve high accuracy. This principle is conceptually related to the “informed training sets” introduced by Wittmann and colleagues^39^ in the context of protein design, which have been shown to provide important advantages in case where data efficiency is a concern. Our obser-vations raise exciting prospects for Design of Experiments strategies purposely aimed at training sequence-to-expression models that are accurate and data-efficient.

Data requirements above 1,000 sequences are still too costly for most practical applications. Further work is thus required on DNA encodings that are maximally informative for training, as well as model architectures that can deliver high accuracy for small datasets. Both strategies have proven highly successful in protein engineering^44,45^, yet their potential for DNA sequence design remains largely untapped. We found that seemingly superficial changes to DNA encodings, e.g. from binary one-hot to ordinal one-hot encodings (Figure 2B), can have substantial impact on predictive performance. Moreover, although biophysical properties such as the CAI or the stability of mRNA secondary structures are not good predictors by themselves^17^, we observed small but encouraging improvements when these were employed in conjunction with one-hot encodings, particularly for small datasets. This suggests that richer mechanistic descriptors, e.g. by including positional information or base-resolution pairing probabilities of secondary structures, may yield further gains in accuracy.

In agreement with other works^46^, we observed that sequence-to-expression models generalize poorly: their accuracy drops significantly for sequences that diverge from those employed for training. This limitation is particularly relevant for strain engineering, where designers may employ predictors to navigate the sequence space beyond the coverage of the training data. A recent study by Vaishnav et al illustrated that these models can indeed generalize well^25^ using a massive training set with over 20,000,000 sequences. Data of such scale are far beyond the capacity of most laboratories, and therefore it appears that poor generalization is likely to become the key limiting factor in the field. We suggest that careful design of training libraries in conjunction with algorithms for controlled sequence design^38^ may help to improve sequence coverage and avoid low-confidence regions of the predictors.

Deep learning models promise to deliver large gains in efficiency across a range of synthetic biology applications. Such models inevitably require training data and there is a risk that the associated experimental costs become an obstacle for many laboratories. In this work we have systematically mapped the relation between data size, diversity and the choice of machine learning models. Our results demonstrate the viability of more data-efficient deep learning models, helping to promote their adoption as a platform technology in microbial engineering.

## IV. METHODS

### A. Data processing

#### Data sources and visualization

The *E. coli* dataset presented by Cambray et al^23^ was obtained from the OpenScience Framework^47^. After removing sequences with missing values for sfGFP fluorescence and growth rate, the dataset contains ~228,000 sequences. In all trained models, we employed the arithmetic mean of sfGFP fluorescence across replicates for the case of normal translational initiation^23^. To visualize sequences in a two dimensional space (Figure 1B), we employed the UMAP algorithm^27^ v0. 5. 1 on sequences featurized on counts of overlapping *k*-mers. We found that the UMAP projection improved for larger *k*, and chose *k* = 4 to achieve a good trade-off between computation time and quality of projection (Supplementary Figure S1); *k*-mer counting was done with custom Python scripts. In all cases, fluorescence measurements were normalized to the maximum sfGFP fluorescence across cells transformed with the same construct averaged over 4 experimental replicates of the whole library^23^.

#### Training, validation, and test data

In Supplementary Figure S3A we illustrate our strategy to partition the full dataset into sets for training, cross-validation and model testing. For each mutational series, we first perform a split retaining 10% of sequences as a fixed held-out set for model testing. We use the remaining sequences as a development set and perform a second split to obtain two partitions for each series. The first partition is for model training and comprises 3200 sequences from which we used varying fractions for training regressors in each series. The second partition was employed for hyperparameter optimization, containing ~400 sequences from each series (10% of the whole series) that we then merged into a large validation set comprising 22,400 sequences (56 series × 400 sequences per series) from all series. We kept the validation set fixed and employed it for hyperparameter optimization of both non-deep and deep models. In all data splits, we stratified the sfGFP fluorescence data to ensure that the phenotype distributions are preserved. Stratification was done with the verstack package, which employs binning for continuous variables; we further customized the code to gain control of the binning resolution.

### B. Model training

#### Non-deep machine learning models

DNA encodings (Table I) were implemented with custom Python code, and all non-deep models were trained using the scikit-learn Python package. To determine model hyperparameters, we used a validation set for all combinations of encodings and regressors. As illustrated in Supplementary Figure S3B, for each model we explored each the hyperparameter search space (Supplementary Table S3) for all encodings using grid search with 10-fold cross validation on 90% of our validation set (~20,000 sequences), using mean squared error (MSE) as performance metric. This resulted in six hyperparameter configurations for each regressor (one for each encoding). For many regressors, we found that the same configuration was optimal for several encodings simultaneously, and we thus settled on most frequent configuration among the six encodings; in case of a tie between configurations, we settled for the one with the best MSE computed on the remaining 10% of our whole validation set.

#### Convolutional neural networks

CNNs were trained on Tesla K80 GPUs from Google Colaboratory^48^. To design the CNN architectures, we use the Sequential class of the Keras package with the TensorFlow backend^49,50^. All CNNs were trained on binary one-hot encoded sequences with mean squared error as loss function, batch size of 64, learning rate 1 × 10^−3^, and using the Adam optimizer^51^. Since ADAM computes adaptive learning rates for each weight of the neural network, we found that the default options were adequate and did not specify a learning rate schedule. We set the maximum number of epochs to 100, and used 15 epochs without loss improvement over the validation set as early stopping criterion to prevent overfitting.

Model hyperparameters were selected with Bayesian optimization implemented in the Hyper-Opt package^35^. Specifically, as shown in Supplementary Figure S3C, we performed five iterations of the HyperOpt routine using 90% of our validation set (~20,000 sequences), where subsets of the search space were evaluated (Supplementary Table S4). We used the Tree of Parzen Estimators (TPE)^52^ as acquisition function, and set the number of architecture combinations to 50. This resulted in five candidate architectures, from which we chose the one with the best validation MSE computed on a stratified sample of size 10% of the whole validation set. The resulting model architecture is described in Supplementary Table S5. To verify that the selected architecture works best for our study, we performed an additional test (Supplementary Figure S9) where we trained CNNs of varying width and depth and compared them to the results in Figure 3C. To achieve this, we perturbed the number of convolutional filters and layers, for width and depth respectively, and trained the resulting architectures using 75% of sequences for each mutational series (Supplementary Figure S9).

#### Model testing

In all cases we did five training repeats on resampled training sets and a fixed test set. Model accuracy was computed as coefficient of determination (*R*^2^) on held-out sequences, averaged across five training repeats. The *R*^2^ score for each training repeat was defined as:

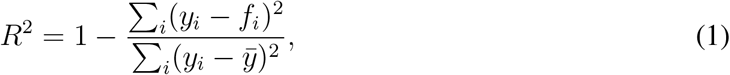

where *y_i_* and *f_i_* are the measured and predicted fluorescence of the *i*^th^ sequence in the test set, I respectively, and *y* is the average fluorescence across the whole test set. Note that for a perfect fit we have *R*^2^ = 1, and conversely *R*^2^ = 0 for baseline model that predicts the average fluorescence (i.e. *f_i_* = *y* for all sequences). Negative *R*^2^ scores thus indicate an inadequate model structure with worse predictions than the baseline model.

### C. Interpretability analysis

For the interpretability results in Figure 3E–G, we employed DeepLIFT^37^ which utilizes back-propagation to produce importance or “attribution” scores for input features, with respect to a baseline reference input. We chose a blank sequence as a reference. We used the *GenomicsDefault* option that implements *Rescale* and *RevealCancel* rules for convolutional and dense layers, respectively. The line plots in Figure 3E are the attribution scores of 30 random test sequences for the CNN and MLP models trained on mutational series 21. The distance heatmaps in Figure 3F were produced by computing the cosine distance between vectors of attribution scores, and then using hierarchical clustering to compare both models. The degree of clustering was quantified by *k*-means scores (Figure 3G); lower scores suggest more clustering of the distance matrix. Results for all other mutational series can be found in S10.

### D. Impact of sequence diversity

#### a. Escherichia coli dataset

The models in Figure 4 were trained on data of constant size and increasing sequence diversity. We successively aggregated fractions of mutational series to create new training sets with improved diversity. We employed the same CNN architecture and training strategy as in Figure 3A with the same hyperparameters (Supplementary Table S5) for all 27 models. To ensure a comparison solely on the basis of diversity, we fixed the size of the training set to 5,800 sequences. To increase diversity, for successive models we sampled training sequences from two additional series, as shown in Figure 4. The specific series for the aggregates were randomly chosen; four training repeats with randomized selection of series can be found in Supplementary Figure S12.

#### b. Saccharomyces cerevisiae dataset

We obtained the promoter dataset presented in Supplementary Figure 4F in Vaishnav et al^25^ from CodeOcean^53^. The data contains 3,929 yeast promoter sequences with YFP fluorescence readouts. To visualize the yeast sequences (Figure 5A), we employed the same strategy as in Figure 1B for the *E. coli* dataset, and used the UMAP algorithm for counts of overlapping 4-mers. Additional details can be found in the Supplementary Text.

For the models in Figure 5B, we first aggregated sequences from the clusters in Figure 5A into twelve groups. We then employed the same strategy as in Figure 4, and successively aggregated fractions of groups to create new training sets with improved diversity. We used the same Random Forest configuration (Supplementary Table S6) for all 5 models. We fixed the size of the training set to 400 sequences, and to increase diversity for successive models, we sampled training sequences from two additional groups at a time (Figure 5B). The specific groups for the aggregates were randomly chosen; four training repeats with randomized selection of groups can be found in Supplementary Figure S13. Additional details can be found in the Supplementary Text.

## Supplementary Text

### Introduction

Here, we detail the promoter dataset in *Saccharomyces cerevisiae* from Vaishnav et al^25^. In the original work, this dataset was employed as a test set to demonstrate the generalization performance of a model trained on ~20M promoter sequences; these results are shown in Supplementary Figure 4F of Vaishnav et al^25^. In our paper, we repurposed this dataset to train sequence-to-expression models with a reduced number of variants. In particular, we employed this new dataset to extend our conclusions on the relation between sequence diversity and model accuracy (Figure 4 in the main text) to a different expression host and a different construct library.

### Dataset

The dataset contains expression levels of 3,929 promoter sequences as measured by a gigantic parallel reporter assay (GPRA)^8^, in which 80bp sequences were embedded within a promoter construct and the expression of a YFP reporter was assayed in a *S. cerevisiae* strain lacking URA3. The library consists of native yeast promoter sequences from 199 genes, each one with an average of 20 random single base mutations. Constructs were cloned within the −160:−80 region, relative to the transcription start site (TSS) of a synthetic promoter scaffold, a critical location for transcription factor binding^54^ and determinant of promoter activity^8^. The promoter construct was placed in a dual reporter plasmid that contains URA3 (used as a selectable marker), a constitutive RFP reported (to control for extrinsic noise), and the YFP reporter under variable control. Finally, yeast cells were cultivated in synthetic defined medium lacking uracil (SD-Ura), sorted into 18 uniformly sized expression bins. Promoters in each bin were sequenced to estimate YFP expression level.

### Impact of sequence diversity on model accuracy

#### Strategy

For the results in Figure 5, we first aggregated the variant clusters into twelve groups, with each group containing variants from ~16 randomly selected clusters. We then trained regressors on group aggregates, in a similar fashion to the analysis in Figure 4. The models in Figure 5 were trained on datasets of constant size and increasing sequence diversity. We successively aggregated fractions of groups to create new training sets with improved diversity.

Given the small size of the training data (~330 sequences/group), we fixed the size of the training set to 400 sequences and focused on training Random Forest models. To increase diversity, for successive models we sampled training sequences from two additional groups, as shown in Figure 5B. The specific groups for the aggregates were randomly chosen; four training repeats with randomized selection of groups can be found in Supplementary Figure S13.

#### Training, validation and test data

To ensure a balanced held-out test set, we uniformly sampled 20% of the 20 sequences that include point mutations for each of the 199 native genes present in the yeast dataset. This resulted in 588 held-out sequences that we reserved for testing all downstream random forest models. The remaining 80% of the total dataset was further partitioned in a similar manner to acquire 780 sequences that we used as a fixed validation set for hyperparameter optimization and 2560 sequences that we used for training.

#### Hyperparameter selection

We performed hyperparameter optimization using binary one-hot encoding and the 780 sequences in the validation set. Hyperparameters were determined via grid search with 10-fold cross-validation. The hyperparameter search space and the resulting random forest configuration, used for all models trained on the yeast dataset, can be found in Supplementary Table S6. We employed the same hyperparameters for all 5 models in Figure 5B in the main text.

## SUPPLEMENTARY FIGURES

**Supplementary Figure S1.**
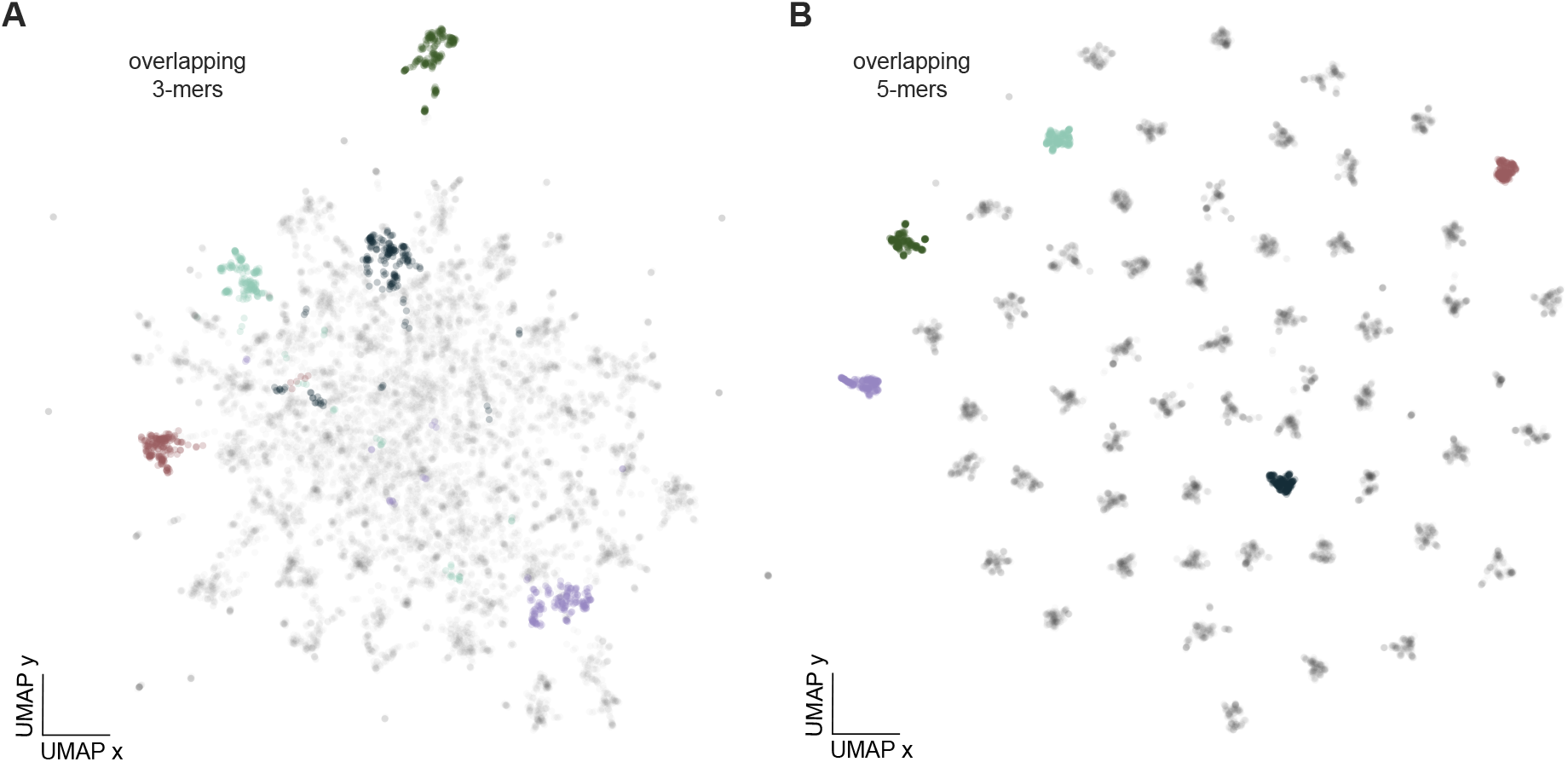
Two dimensional projections of the 228,000 sequences employed for training. **(A)** UMAP projection for overlapping 3-mers; the choice of 3-mers does not have enough granularity for UMAP to distinguish between mutational series. **(B)** UMAP projection for overlapping 5-mers, which show a similar cluster structure as the one computed with 4-mers in Figure 1B. Colour coding is the same as in Figure 1B.

**Supplementary Figure S2.**
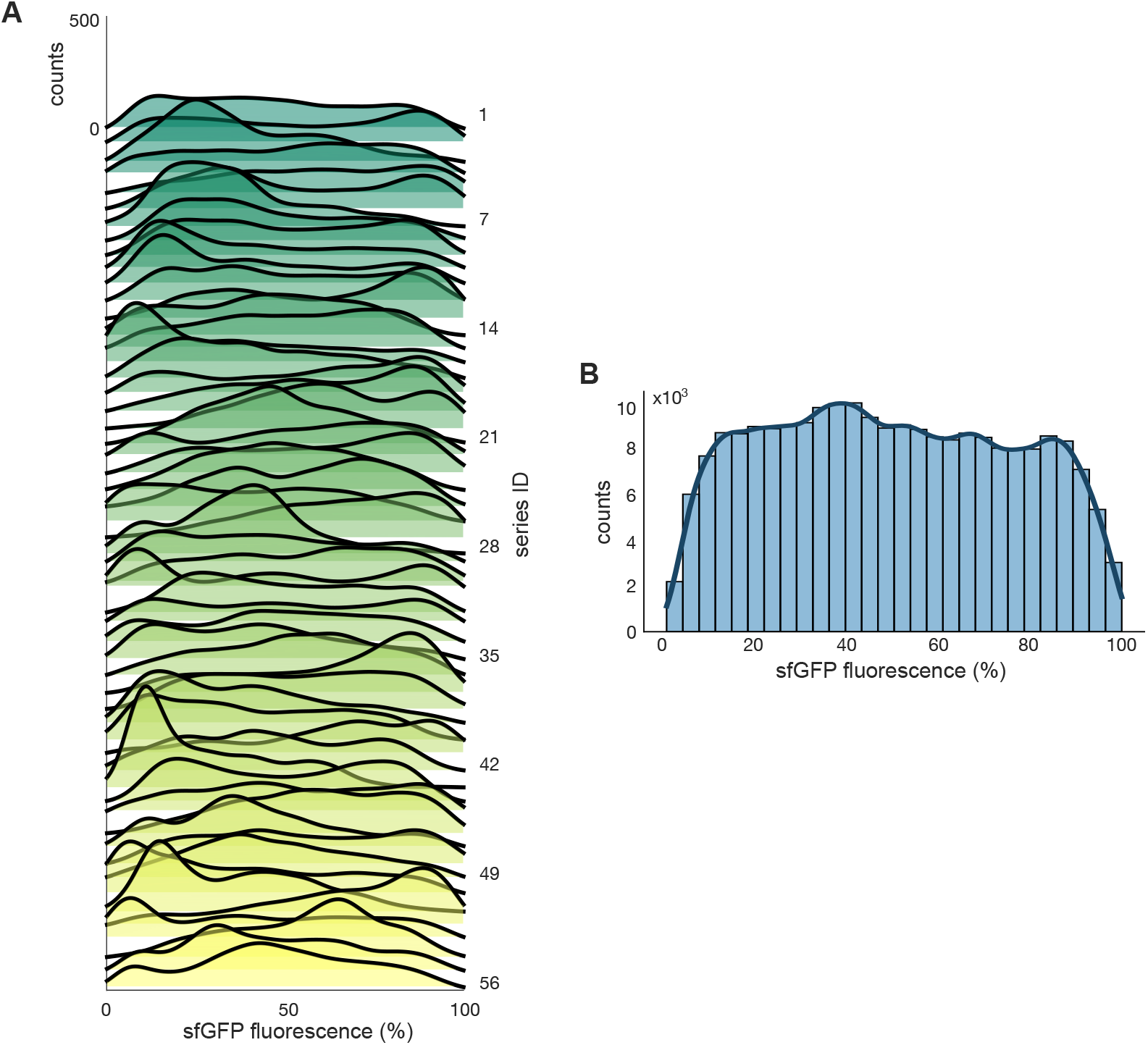
Distributions of sfGFP fluorescence from the dataset in Cambray et al^23^. **(A)** Phenotypic distributions for each of the 56 mutational series. **(B)** Phenotypic distribution of the complete dataset with 56 mutational series and ~228,000 sequence variants. Shown distributions are Gaussian kernel density estimates of fluorescence measurements averaged across four experimental replicates. Measurements are normalized to the maximum sfGFP fluorescence in the whole library.

**Supplementary Figure S3.**
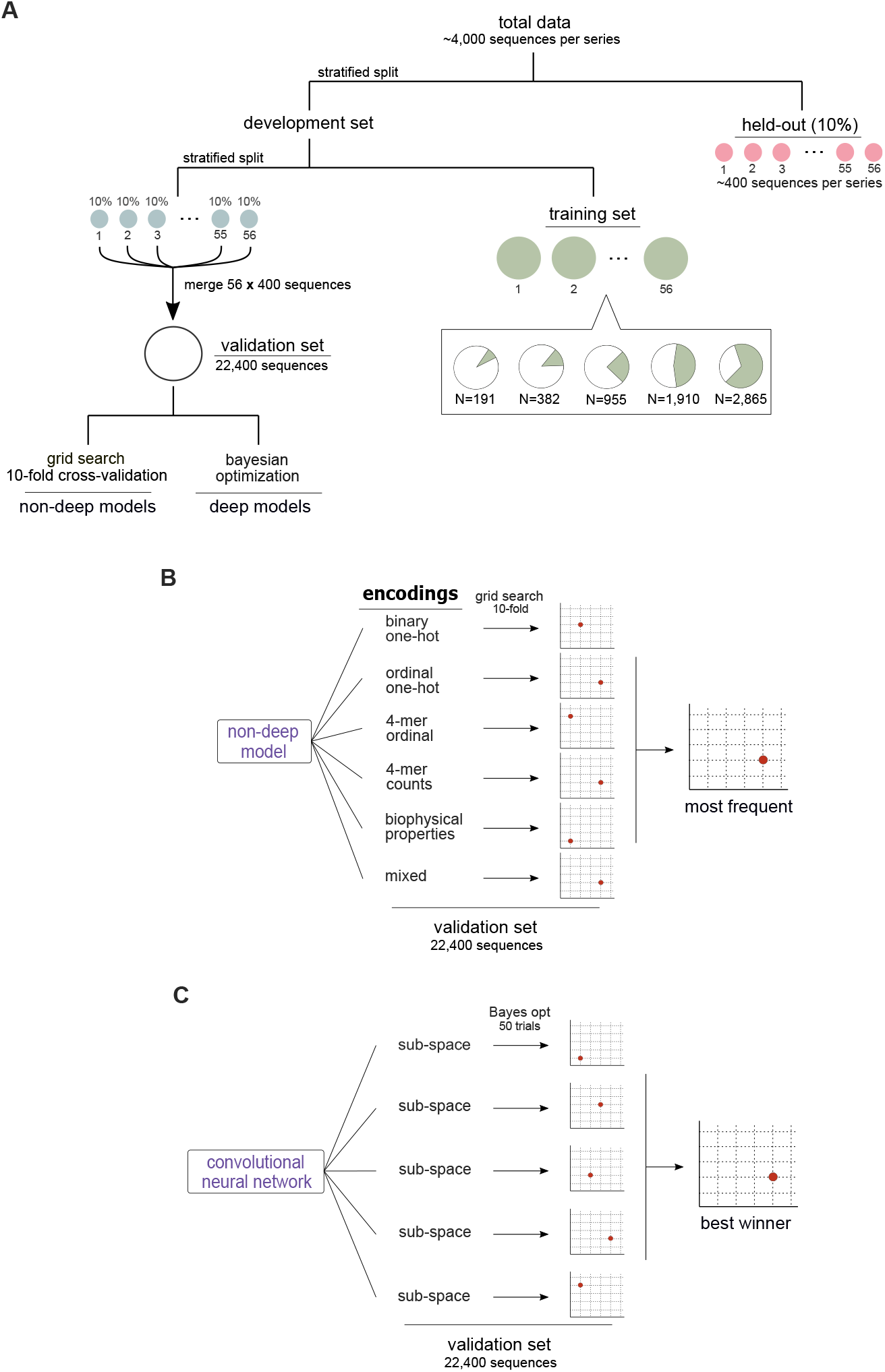
Data splitting and hyperparameter optimization strategy. **(A)** Schematic of data partitioning into separate sets for training, validation and testing. We first held-out 10% of each series (~400 sequences) for model testing; none of these sequences were used for training or cross-validation. The remaining sequences were further partitioned into two: a training set, which we employed to train models on varying data sizes, and a large validation set with 22,400 sequences drawn from all mutational series. The validation set was employed to determine model hyperparameters with 10-fold cross-validation (non-deep models) and Bayesian optimization (deep models). **(B)** Hyperparameter tuning for non-deep models. We explored the hyperparameter space for each regressor (Supplementary Table S2) using grid search and 10-fold cross-validation for each DNA encoding on 90% of the full validation set. This resulted in one configuration per encoding, from which we selected the most frequent configuration among the six encodings. For cases where there was no single most frequent configuration, we selected the one with the smallest mean squared error (MSE) on the remaining 10% of the validation set. **(C)** Hyperparameter tuning for CNN models. We ran five iterations of Bayesian optimization implemented in the HyperOpt package (Supplementary Table S4) and obtained five candidate architectures, from which we settled on the one with smallest MSE on the remaining 10% of the validation set.

**Supplementary Figure S4.**
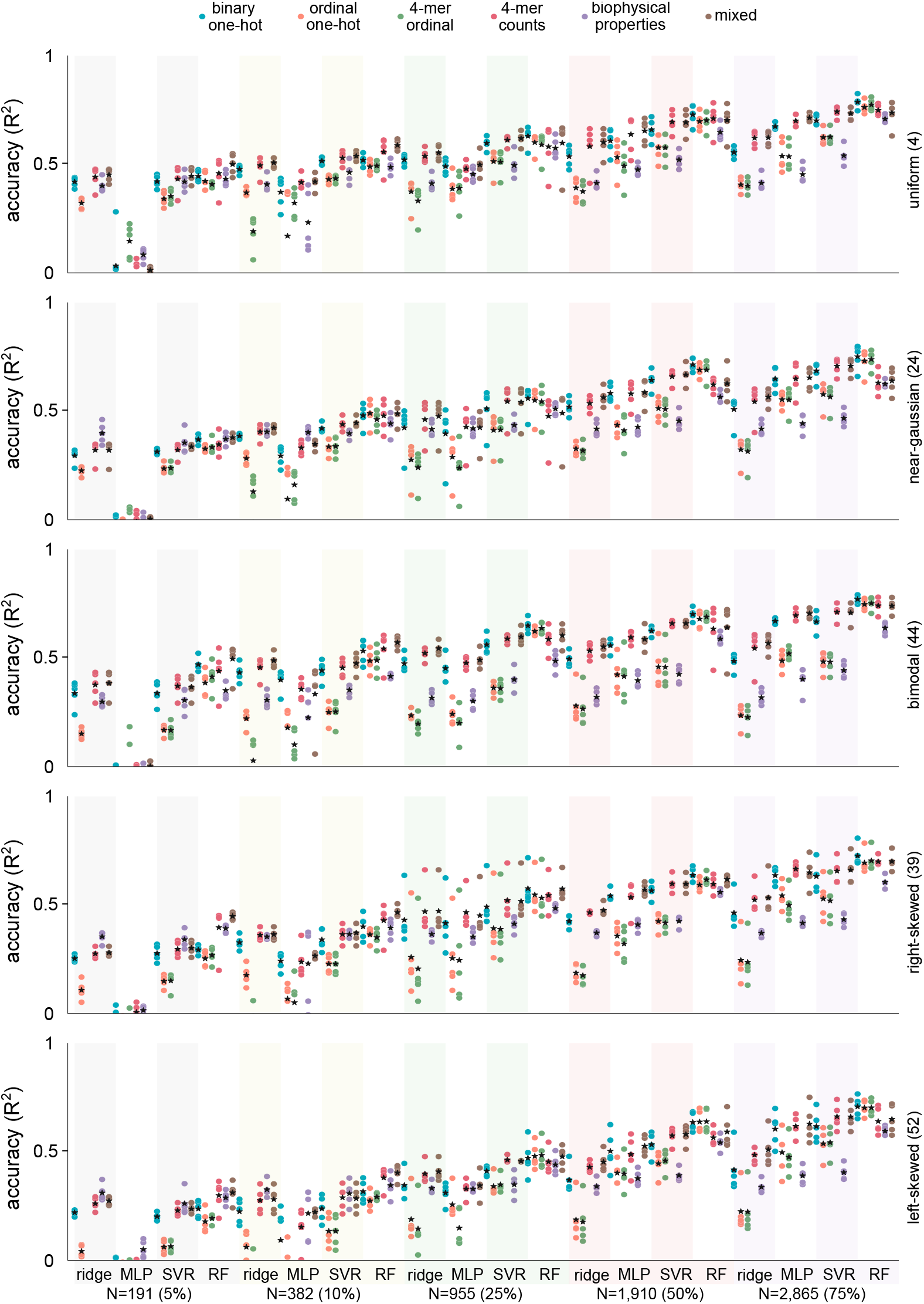
Cross-validation of non-deep models in Figure 2B. Shown are the *R*^2^ scores computed between measured and predicted fluorescence in held-out sets of sequences; dots are the *R*^2^ scores for each training repeat and stars denote the mean accuracy across the five training repeats (Monte Carlo cross-validation); the plots show 600 models in total (i.e. 4 regressors ×5 data sizes ×6 encodings ×5 mutational series). In each training repeat, we held-out 10% of randomly chosen variants in each mutational series, and trained all models on the specified number of samples (N); note that in each training repeat, the test set was kept constant for all models to ensure fair comparisons across models, i.e. all models were tested on the same set of held-out sequences. Overall, the results show robust accuracy across cross-validation runs, particularly for the high-accuracy models. There is some variation in *R*^2^ values for low and mid accuracy regressors, but the best performing models show little evidence of overfitting.

**Supplementary Figure S5.**
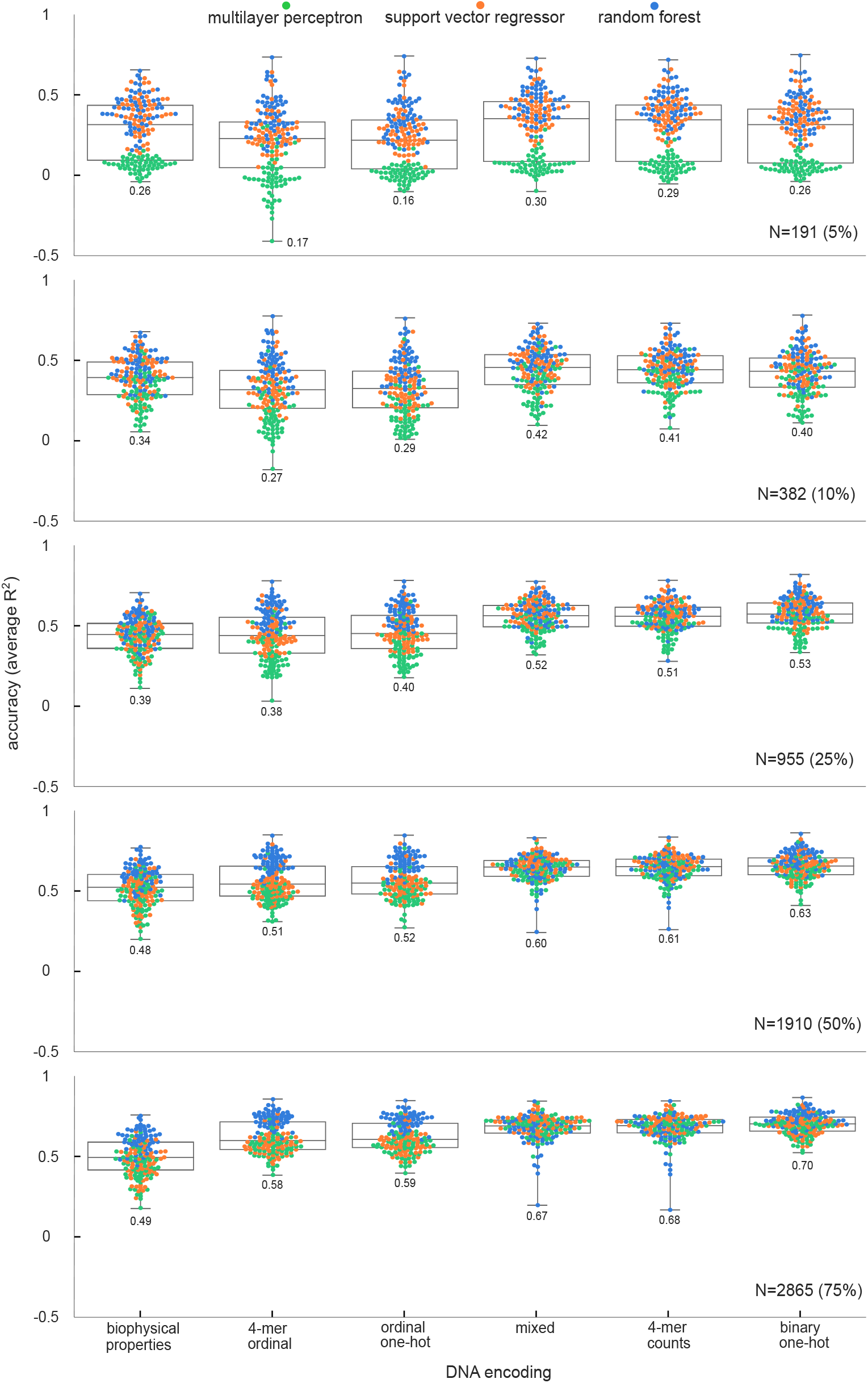
Accuracy of non-deep models trained on the whole sequence space. We trained 5,040 models (3 regressors × 5 data sizes × 6 encodings × 56 mutational series); we excluded the ridge regressor due to its poor performance (Figure 2B). Dots in the swarm plots are the prediction accuracy scores for each model, computed as the **R*^2^* on a fixed held-out dataset with 10% of sequences of each series, and averaged across 5 training repeats. Random forests with binary one-hot encoding provide the best accuracy and the least sensitivity to the shape of the sequence space. Binary one-hot encoding also lead to more consistent accuracy across series, as reflected by narrower distributions of *R*^2^ values.

**Supplementary Figure S6.**
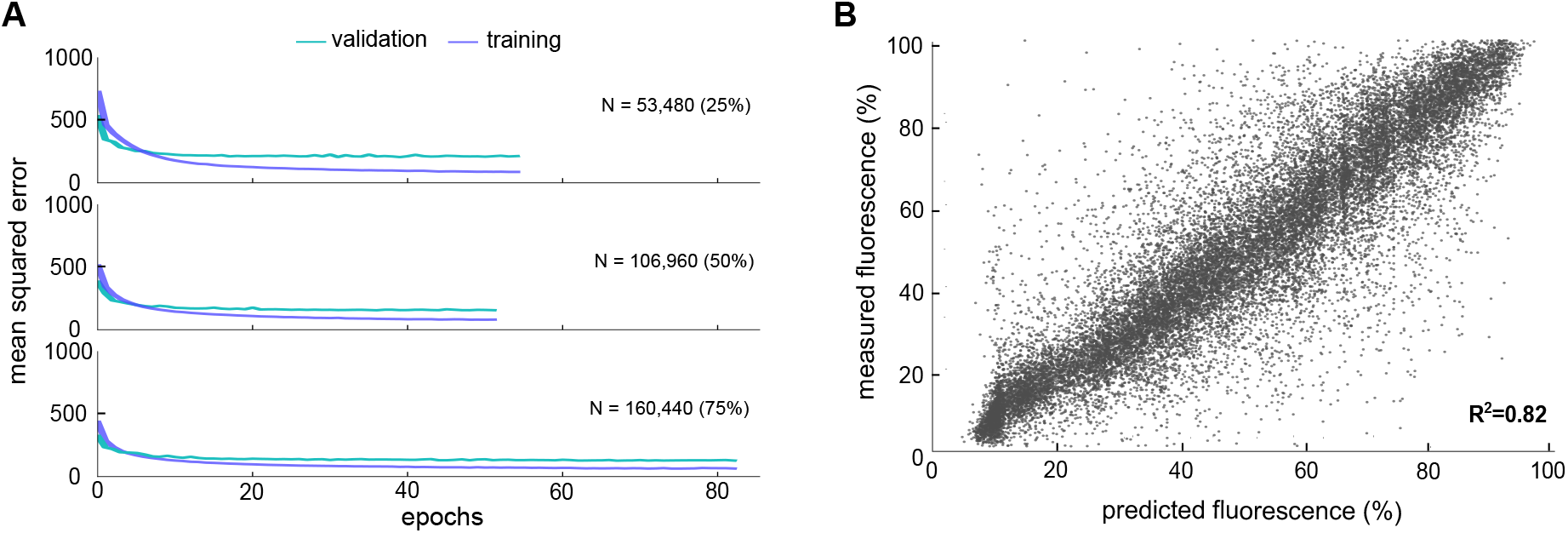
Convolutional neural network trained on all mutational series. **(A)** Learning curves computed as the mean squared error against training epoch for the validation (cyan) and training (purple) sets for CNNs trained on 25%, 50%, and 75% of all sequences (Figure 3B). In all cases, we use the same validation set, containing 22,400 sequences aggregated over all 56 mutational series (Supplementary Figure S3A), and use 15 epochs without loss improvement on the validation set as early stopping criterion to prevent overfitting. **(B)** Predictions of the CNN from Figure 3B trained on 75% of all sequences and evaluated on held-out sequences (10% of total sequences). We note that although *R*^2^= 0.82 is comparable to the random forests models in Figure 2B, those models were trained and tested on a single mutational series; the CNN produces accurate predictions across all mutational series.

**Supplementary Figure S7.**
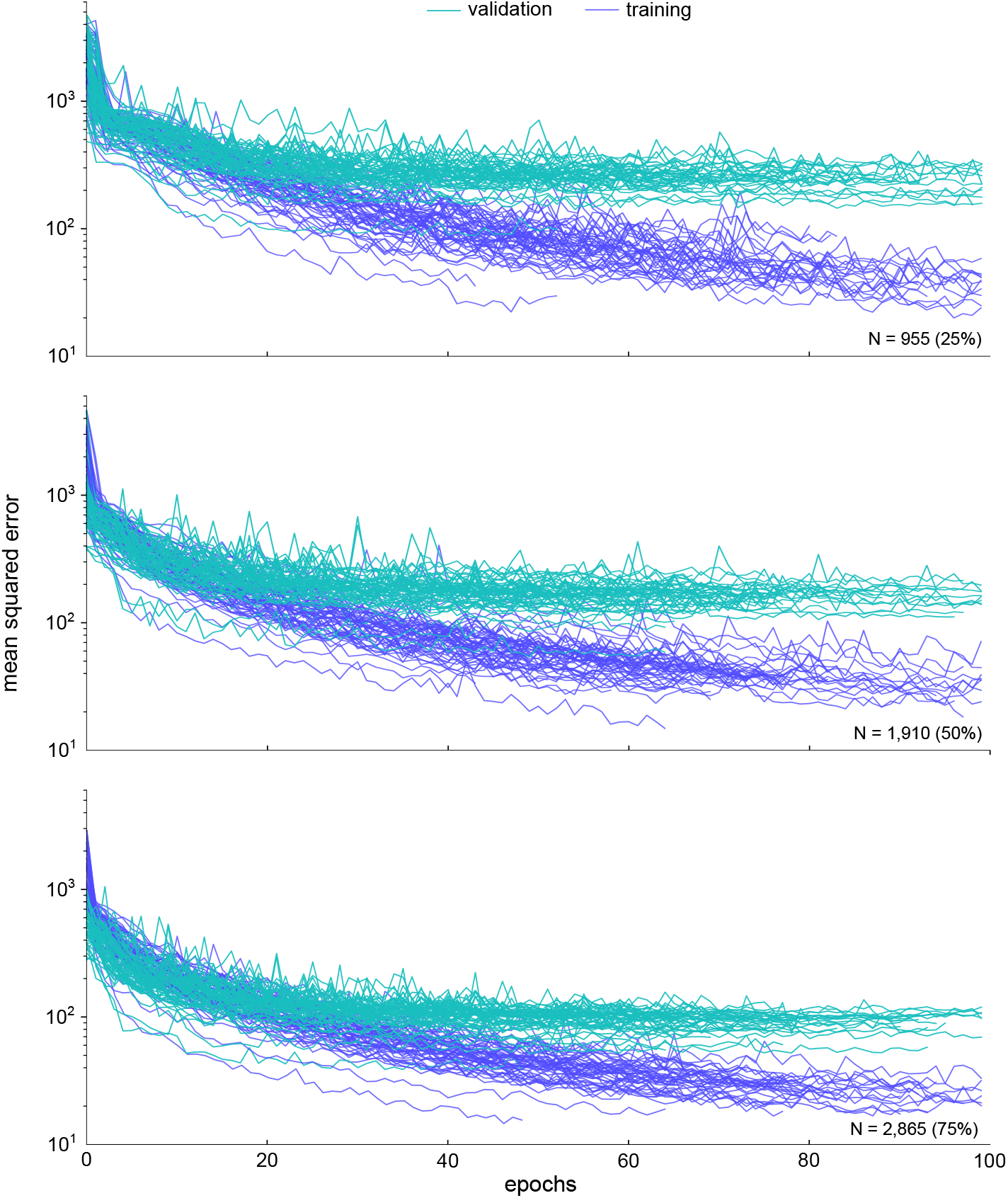
Learning curves for convolutional neural networks trained on each mutational series. Plots show the validation (cyan) and training (purple) mean squared error against training epochs for individual CNNs trained on 25%, 50%, and 75% of the sequences in each mutational series. We use fixed validation sets with 10% of sequences from each series, and 15 epochs without loss improvement over the validation set as early stopping criterion to prevent overfitting.

**Supplementary Figure S8.**
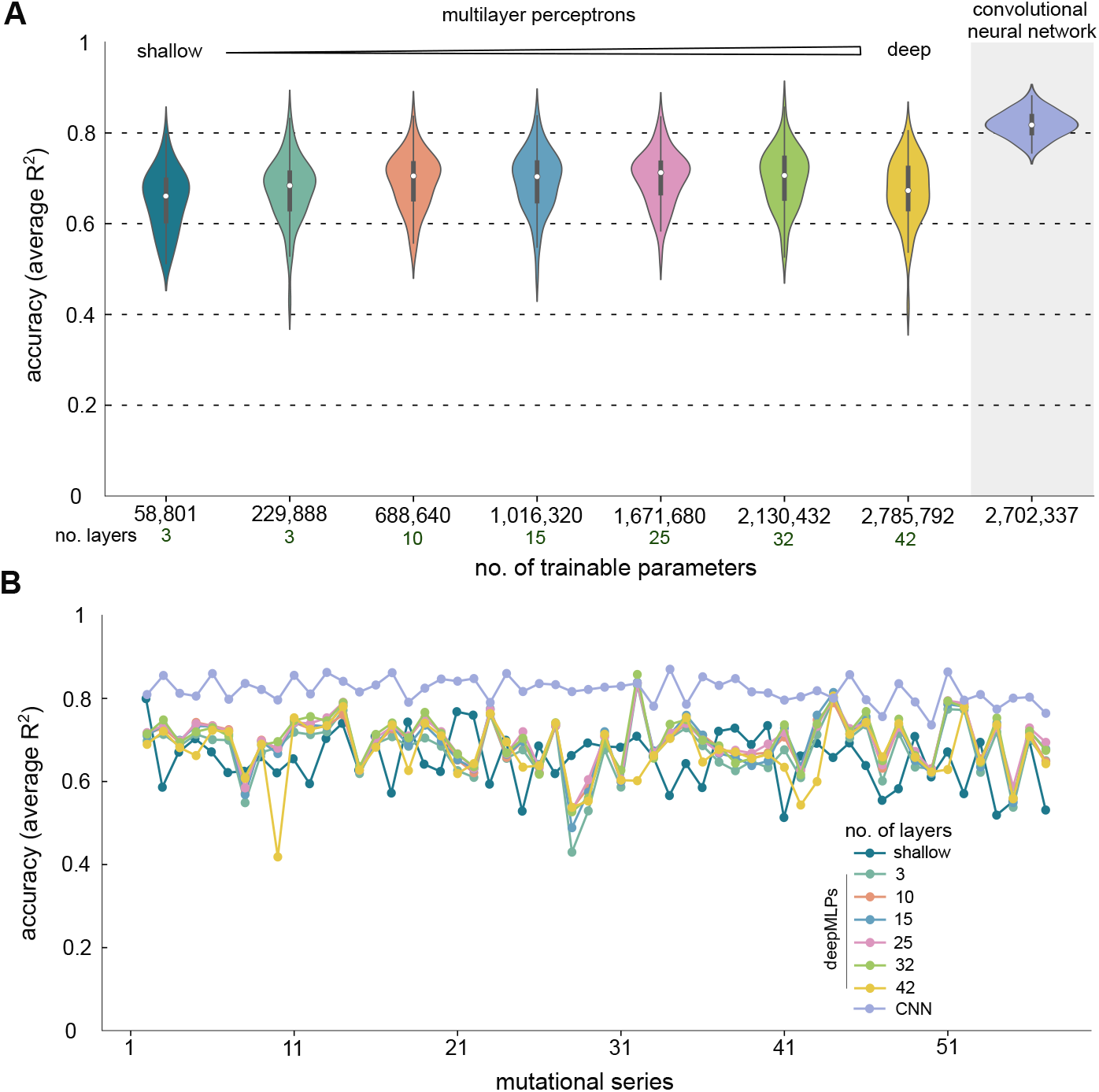
Performance comparison between deep MLPs and CNNs. (**A**) Prediction accuracy of MLPs of increasing depth against the CNNs in Figure 3C for each of the 56 mutational series using binary one-hot encoding and 75% of sequences for training. The shallow and the first deep MLP have the same number of layers, but different number of neurons per layer, with 100 versus 256 neurons respectively. The remaining deep MLPs contain 256 neurons per layer, and were implemented in scikit-learn. The CNNs outperformed the deep MLPs even in cases when they both have a comparable number of trainable parameters (>2.7M parameters). This suggests that the convolutional layers can extract sequence features that are highly informative for regressing the protein expression level. Violin plots show the distribution of the 56 *R*^2^ scores for each model averaged across 5 training repeats; *R*^2^ values were computed on held-out sequences (10% of sequences per series). Deep MLPs were trained with the ReLU activation function and mean squared error as loss function, learning rate 1 × 10^-3^, and using the Adam optimizer^51^. To prevent overfitting, we set the maximum number of epochs to 120 and used 15 epochs without loss improvement over the validation set as early stopping criterion. **(B)** *R*^2^ scores averaged across five training repeats for each model in panel A. The deep MLPs marginally outperform the CNNs in only two mutational series (no. 32 and 43).

**Supplementary Figure S9.**
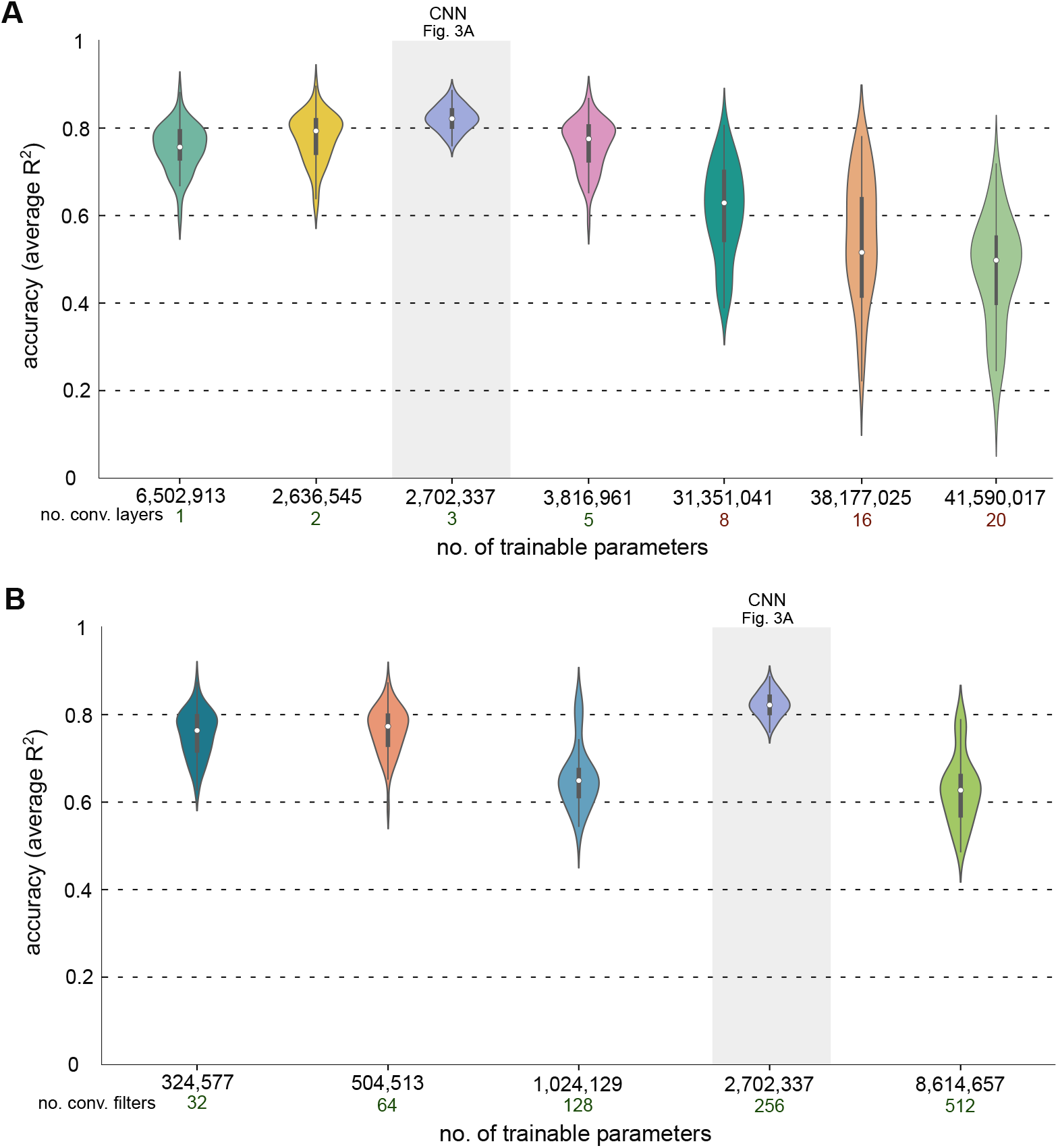
Perturbation analysis of CNNs in Figure 3A. (**A**) Prediction accuracy of retrained CNNs with a variable number of convolutional layers, against the CNN in Figure 3A for each of the 56 mutational series using binary one-hot encoding and 75% of sequences for training. We varied the number of convolutional layers from one to twenty and froze all other hyperparameters to the values in Supplementary Table S4. Violin plots show the distribution of the 56 *R*^2^ scores for each mutational series computed on held-out sequences (10% of sequences per series) and averaged across 5 training repeats; for the deeper CNNs with {8,16, 20}convolutional layers we used only one training repeat. The CNN with one convolutional layer has a larger number of trainable parameters because of the lack of a max pooling layer. In CNN architectures with few convolutional layers, most of the parameters are concentrated in the dense layers. After flattening, the inputs to the dense layers are higher-dimensional than for deeper CNNs. Note that for the deeper CNNs with {8, 16, 20}convolutional layers, we removed max pooling layers and included batch normalization to stabilize the learning process. **(B)** Prediction accuracy of CNNs with varying width (i.e. number of filters per layer) against the CNN in Figure 3A for each of the 56 mutational series using binary one-hot encoding and 75% of sequences for training. To implement CNNs of varying widths, we retrained CNNs with a variable number of convolutional filters in the three convolutional layers; all other CNN hyperparameters were frozen to the values in Supplementary Table S4. In line with expectation, in panels A and B we found that the Bayesian optimized architecture (gray band, shown in Figure 3A) outperforms the other architectures. In panel B, violin plots show the distribution of the 56 *R*^2^ scores for each mutational series computed on held-out sequences (10% of sequences per series) and averaged across 5 training repeats.

**Supplementary Figure S10.**
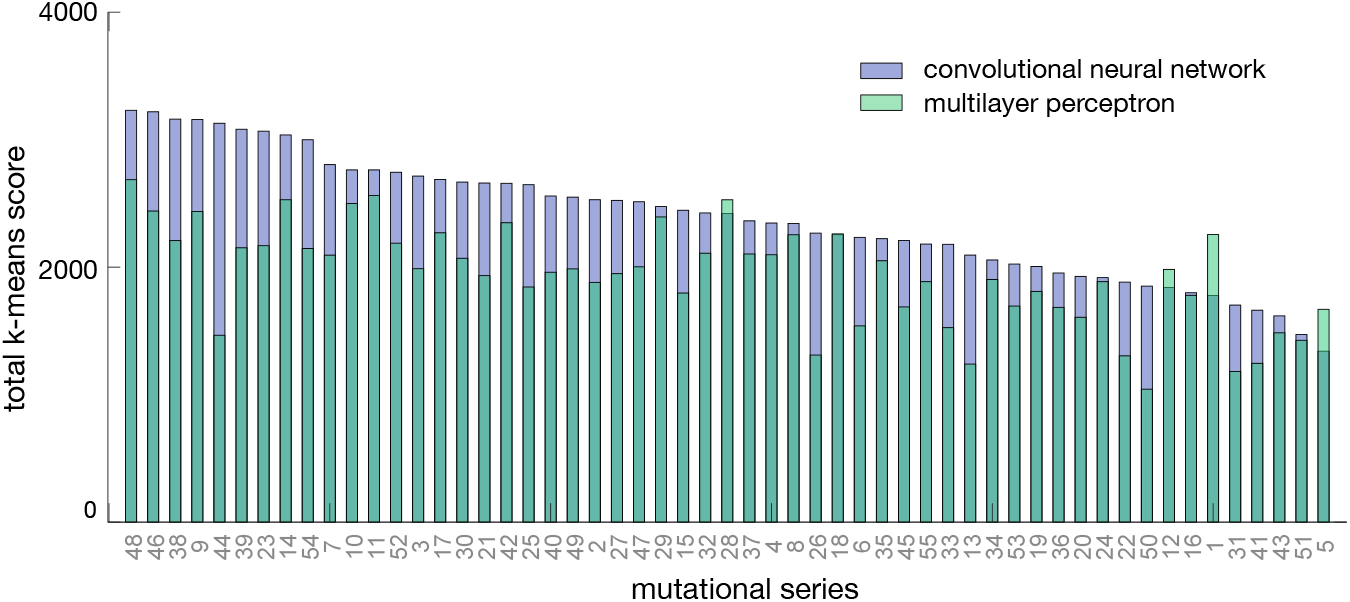
Comparison of neural networks using DeepLIFT scores. We performed *k*-means clustering on the attribution distance matrices (Figure 3F) for each of the 56 models. Bars show the total *k*-means score for test sets in each of the CNNs shown in Figure 3C, averaged across 20 runs of the *k*-means clustering algorithm; the total score is defined as 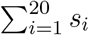 where *s_i_* is the clustering score for a fixed number of clusters *k*. Clusters were computed using DeepLIFT^37^ scores as feature vectors. The CNNs display higher *k*-means scores in all but four mutational series, which suggests that they are better at discriminating between similar sequences in a test set; all test sets contain 10% of sequences of each mutational series (~450 sequences/series).

**Supplementary Figure S11.**
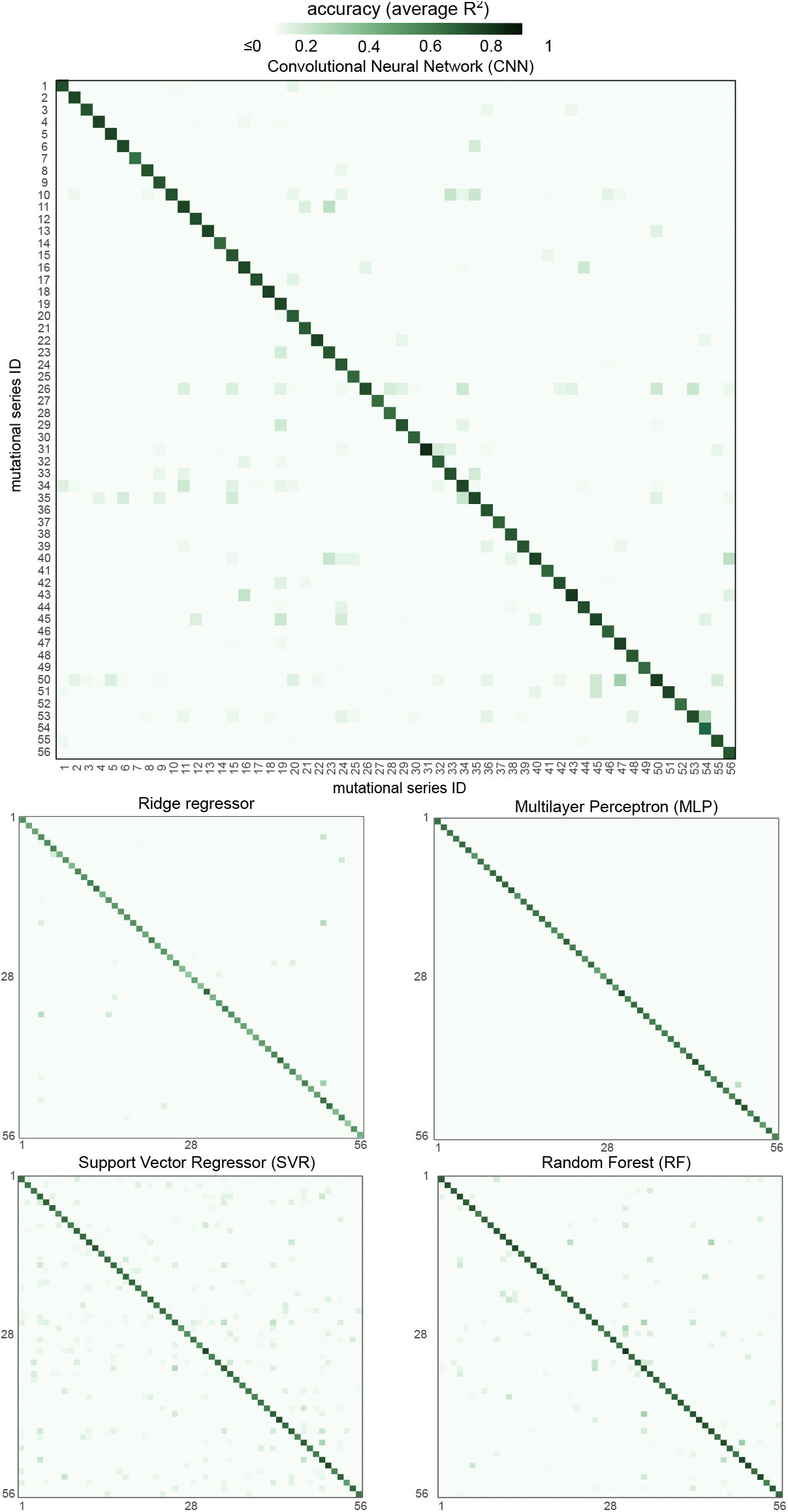
Generalization performance of machine learning models trained on a single mutational series. Heatmaps show the accuracy (*R*^2^) of CNNs trained on a single series and tested in all other series. Models were trained on 75% of a single mutational series and tested on held-out sequences from every other series (10% of each series). Values in the diagonal are the accuracy when tested on 10% of held-out sequences from the same series employed for training. Accuracy is reported as the *R*^2^ computed on a held-out test set and averaged across five training repeats. The results indicate that model generalization is extremely poor, with all models achieving low or negative cross-series accuracy.

**Supplementary Figure S12.**
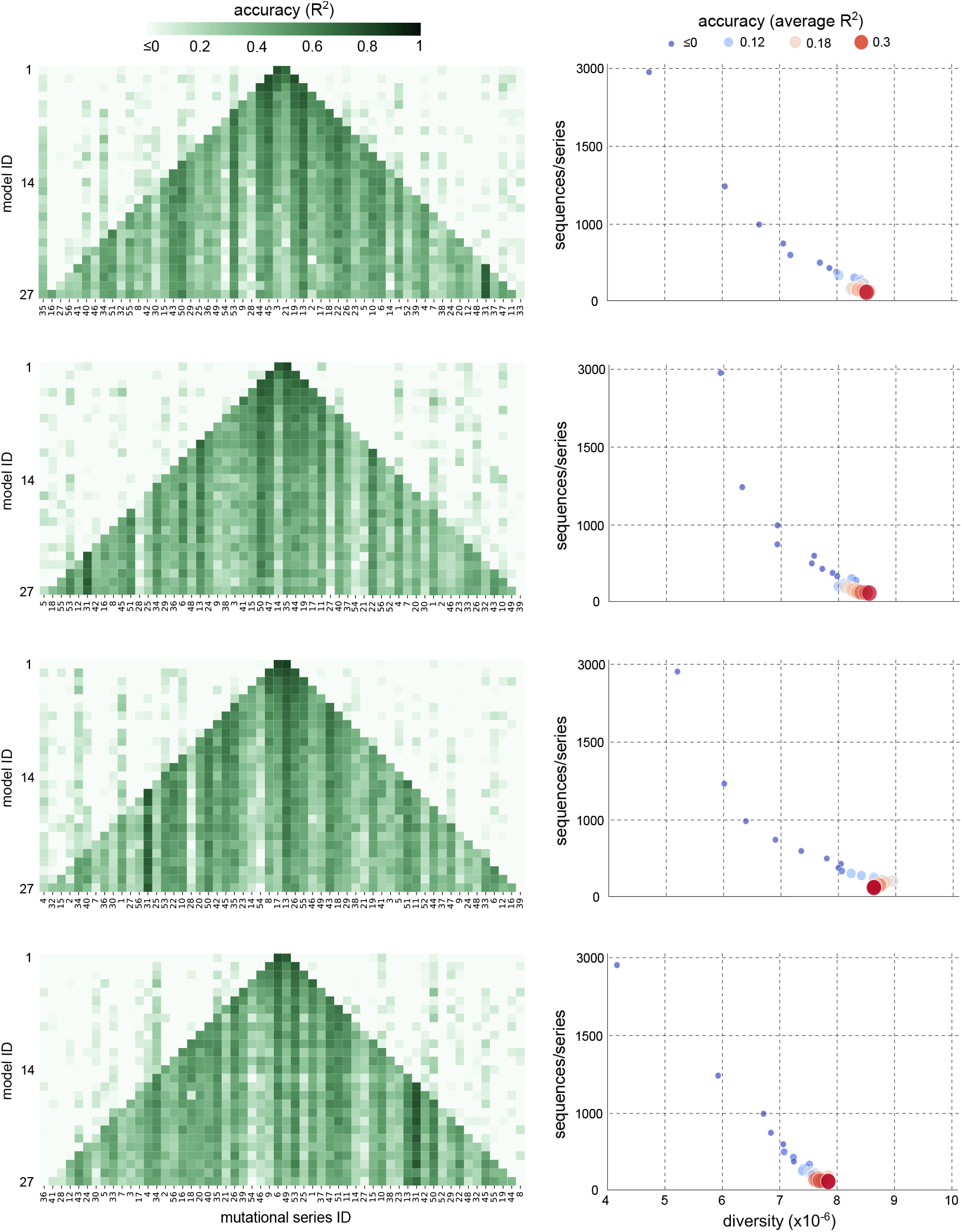
Convolutional neural networks trained on an increasingly diverse sequence space. Shown are four repeats of the computational experiment shown in Figure 4, with randomized selection of mutational series employed for training.

**Supplementary Figure S13.**
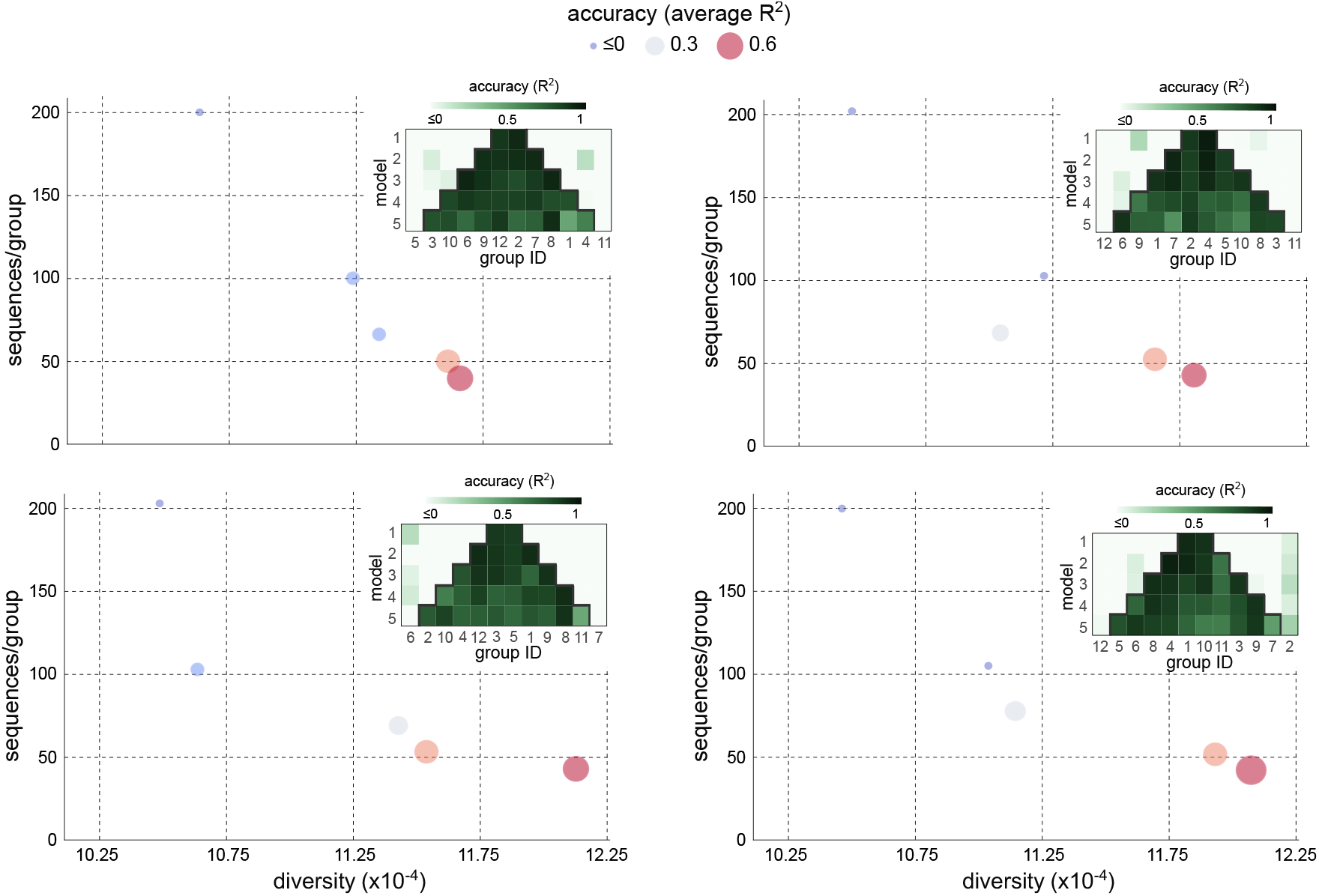
Random forests trained on an increasingly diverse sequence space for *Saccharomyces cerevisiae*. Shown are four repeats of the computational experiment shown in Figure 5, with randomized selection of groups employed for training.

## SUPPLEMENTARY TABLES

**Supplementary Table S1.**
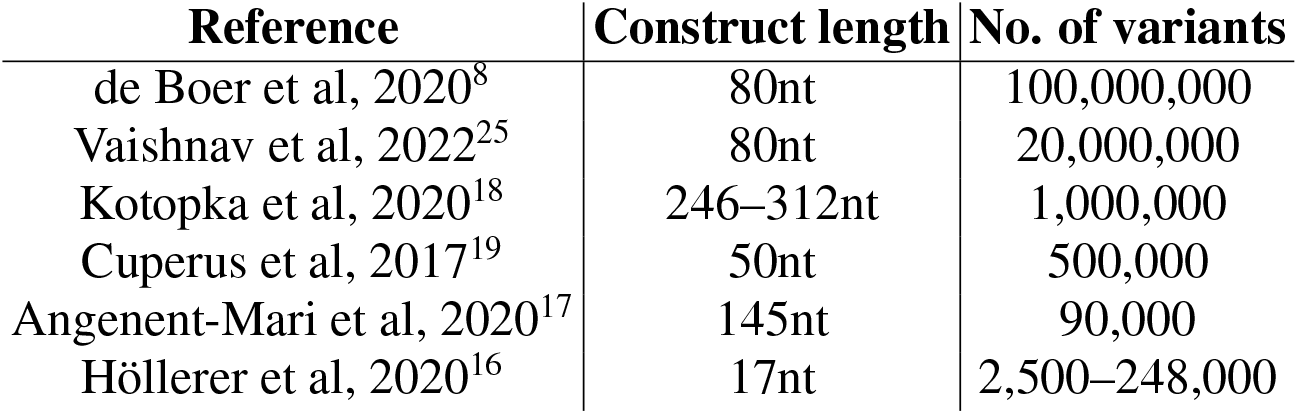
Recent sequence-to-expression machine learning models. The list focuses on studies on prediction from short DNA or RNA sequences. The list excludes studies focused on whole gene prediction^14^ or those that focus on other phenotypes beyond protein expression, such as transcription factor binding^12^ or mRNA levels^55^; data sizes have been rounded.

**Supplementary Table S2.** Search space for hyperparameters for non-deep models in Figure 2. We performed an exhaustive grid-search with 10-fold cross-validation over the specified parameter values of each regressor, for all encodings. Square brackets correspond to ranges where all integer values were used; we report the increment value for cases where only parts of the range were used. Curly brackets specify the set of values used.

| **Regressor** | **Hyperparameter** | **Value** |
| --- | --- | --- |
| support vector regressor | regularization ($C$) | $[1, 50]$ |
| | margin tolerance ($\epsilon$) | $[0.1, 2]$ |
| multilayer perceptron | activation function | $\{\text{ReLU}, \text{tanH}\}$ |
| | hidden layers | $[1, 5]$ |
| | no. of neurons | $[100, 400]$ incr. of 50 |
| random forest | no. of estimators | $[5, 100]$ incr. of 10 |
| | maximum depth | $[15, 100]$ incr. of 5 |
| | min samples per leaf | $[1, 12]$ |
| | min samples to split | $[2, 12]$ |

**Supplementary Table S3.** Hyperparameters for non-deep machine learning regressors. We employed the same hyperparameters for all combinations of mutational series and DNA encodings in all models, except the ridge regressor. The regularization strength of the ridge regressor was optimized on a case-by-case basis in the range shown.

| **Regressor** | **Hyperparameter** | **Value** |
| --- | --- | --- |
| ridge regressor | regularization ($\alpha$) | $[10^{-1}, 10^2]$ |
| support vector regressor | kernel method | RBF |
| | regularization ($C$) | 30 |
| | margin tolerance ($\epsilon$) | 0.5 |
| multilayer perceptron | activation function | ReLU |
|  | hidden layers | 3 |
|  | no. of neurons | 100 |
| random forest | no. of estimators | 25 |
|  | maximum depth | 30 |
|  | min samples per leaf | 3 |
|  | min samples to split | 2 |

**Supplementary Table S4.**
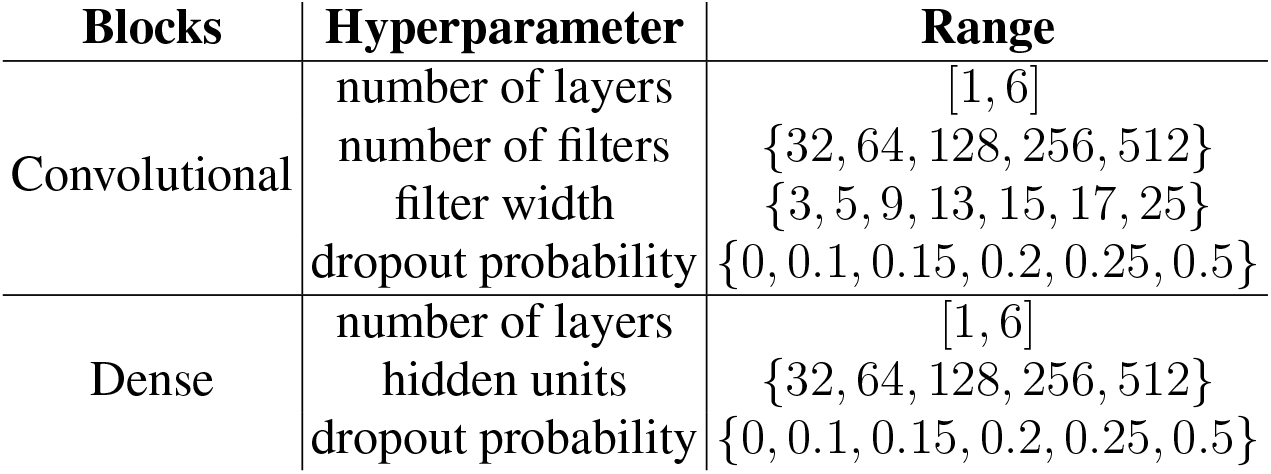
Search space for hyperparameters of the convolutional neural network in Figure 3A. We used subsets of the search space to run five iterations of the HyperOpt routine. In each run, HyperOpt performs Bayesian optimisation and assesses 50 combinations of hyperparameters in an informed manner. Square brackets correspond to ranges where all integer values were used as input, whereas for curly brackets only the specified values were used.

| **Blocks** | **Hyperparameter** | **Range** |
| --- | --- | --- |
| Convolutional | number of layers | [1, 6] |
|  | number of filters | {32, 64, 128, 256, 512} |
|  | filter width | {3, 5, 9, 13, 15, 17, 25} |
|  | dropout probability | {0, 0.1, 0.15, 0.2, 0.25, 0.5} |
| Dense | number of layers | [1, 6] |
|  | hidden units | {32, 64, 128, 256, 512} |
|  | dropout probability | {0, 0.1, 0.15, 0.2, 0.25, 0.5} |

**Supplementary Table S5.** Architecture of the convolutional neural network. For fair comparisons across datasets, we used the same architecture and hyperparameters in all CNNs. We employed 2D convolutions, without skip connections, and set the padding option to *same* for all layers, to ensure that all parts of the sequences equally employed for training. We also included a max pooling layer to reduce the number of trainable parameters.

| **Blocks** | **Hyperparameter** | **Value** |
| --- | --- | --- |
| Convolutional (1-3) | number of filters | 256 |
|  | filter width | 13 |
|  | dropout prob. | 0.15 |
|  | activation | ReLU |
|  | max-pooling | (2,2) |
| Dense (4-7) | hidden units | 256 |
|  | dropout prob. | 0.1 |
|  | activation | ReLU |
| Dense (final) | unit | 1 |

**Supplementary Table S6.** Hyperparameter tuning for random forest regressor trained on S. *cerevisiae* promoter data^25^. We used the same search space for the random forest as in Supplementary Table S2 and performed an exhaustive grid-search with 10-fold cross-validation with one-hot encoding. Square brackets correspond to ranges where all integer values were used; we report the increment value for cases where only parts of the range were used. Curly brackets specify the set of values used. We employed the same hyperparameter values for all random forest models in Figure 5B.

| Hyperparameter | Search space | Chosen value |
| --- | --- | --- |
| number of estimators | [5, 100] incr. of 10 | 50 |
| maximum depth | [15, 100] incr. of 5 | 30 |
| min samples per leaf | [1, 12] | 3 |
| min samples to split | [2, 12] | 4 |

## Notes

### Competing Interest Statement

The authors have declared no competing interest.

### Summary of Updates

Updated text and figures

